# TIGAR: An Improved Bayesian Tool for Transcriptomic Data Imputation Enhances Gene Mapping of Complex Traits

**DOI:** 10.1101/507525

**Authors:** Sini Nagpal, Xiaoran Meng, Michael P. Epstein, Lam C. Tsoi, Matthew Patrick, Greg Gibson, Philip L. De Jager, David A. Bennett, Aliza P. Wingo, Thomas S. Wingo, Jingjing Yang

**Affiliations:** School of Biology, Georgia Institute of Technology, Atlanta, GA, 30332, USA; Department of Biostatistics and Bioinformatics, Emory University School of Public Health, Atlanta, GA, 30322, USA; Center for Computational and Quantitative Genetics, Department of Human Genetics, Emory University School of Medicine, Atlanta, GA, 30322, USA; Department of Dermatology; Department of Computational Medicine & Bioinformatics; Department of Biostatistics, University of Michigan, Ann Arbor, MI, 48109, USA; Department of Dermatology, University of Michigan Medical School, Ann Arbor, MI, 48109, USA; Medical Center Neurological Institute, Columbia University, New York, NY, 10032, USA; Rush Alzheimer’s Disease Center, Rush University Medical Center, Chicago, IL, 60612, USA; Division of Mental Health, Atlanta VA Medical Center, Decatur, GA, USA; Department of Psychiatry and Behavioral Sciences, Emory University School of Medicine, Atlanta, GA, 30322, USA; Department of Neurology, Emory University School of Medicine, Atlanta, GA, 30322, USA

## Abstract

The transcriptome-wide association studies (TWAS) that test for association between the study trait and the imputed gene expression levels from cis-acting expression quantitative trait loci (cis-eQTL) genotypes have successfully enhanced the discovery of genetic risk loci for complex traits. By using the gene expression imputation models fitted from reference datasets that have both genetic and transcriptomic data, TWAS facilitates gene-based tests with GWAS data while accounting for the reference transcriptomic data. The existing TWAS tools like PrediXcan and FUSION use parametric imputation models that have limitations for modeling the complex genetic architecture of transcriptomic data. Therefore, to improve on this, we propose to use a Bayesian method that assumes a data-driven nonparametric prior to impute gene expression. The nonparametric Bayesian method is flexible and general because it includes both of the parametric imputation models used by PrediXcan and FUSION as special cases. Our simulation studies showed that the nonparametric Bayesian model improved both imputation *R*^2^ for transcriptomic data and the TWAS power over PrediXcan. In real applications, our nonparametric Bayesian method fitted transcriptomic imputation models for 57.6% more genes over PrediXcan, thus improving the power of follow-up TWAS. Hence, the nonparametric Bayesian model is preferred for modeling the complex genetic architecture of transcriptomes and is expected to enhance transcriptome-integrated genetic association studies. We implement our Bayesian approach in a convenient software tool “**TIGAR**” (**T**ranscriptome-**I**ntegrated **G**enetic **A**ssociation **R**esource), which imputes transcriptomic data and performs subsequent TWAS using individual-level or summary-level GWAS data.

## Introduction

Genome-wide association studies (GWAS) have successfully identified thousands of genetic risk loci for complex traits. However, the majority of these loci are located within noncoding regions whose molecular mechanisms remain unknown^1–3^. Recent studies have shown that these associated regions were enriched for regulatory elements such as enhancers (H3K27ac marks)^4; 5^ and expression of quantitative trait loci (eQTL)^6; 7^, suggesting that the genetically regulated gene expression might play a key role in the biological mechanisms of complex traits. Multiple studies have recently generated rich transcriptomic datasets for diverse tissues of the human body, e.g., the Genotype-Tissue Expression (GTEx) project for 44 human tissues^6^, Genetic European Variation in Health and Disease (GEUVADIS) for lymphoblastoid cell lines^8^, Depression Genes and Networks (DGN) for whole-blood samples^9^, and the North American Brain Expression Consortium (NABEC) for cortex tissues^10^. Previous studies^11–16^ have also shown that integrating transcriptomic data in GWAS can help identify novel functional loci.

The majority of GWAS projects do not possess transcriptomic data and thus cannot directly conduct integrative analysis. Existing studies^11; 12^ have shown that one can impute the genetically regulated gene expression (GReX) within such GWAS projects by using reference datasets like GTEx^6^ and GEUVADIS^8^, and then test for association between imputed GReX for GWAS samples and the trait of interest — referred to as transcriptome-wide association studies (TWAS)^11; 12^. Specifically, the gene expression imputation models are fitted by regressing assayed gene-expression levels on cis-eQTL genotypes with reference dataset. For examples, the PrediXcan^11^ method uses an Elastic-Net^17^ variable selection model on reference dataset to estimate the cis-eQTL effect-sizes that are used to impute the GReX for GWAS samples, while the FUSION^12^ tool implements a Bayesian sparse linear mixed model (BSLMM)^18^ to estimate the cis-eQTL effect-sizes.

Basically, the Elastic-Net^17^ model used by PrediXcan^11^ assumes a combination of LASSO^19^ (*L*_1_) and Ridge^20^ (*L*_2_) penalties on the cis-eQTL effect-sizes, which is equivalent to a Bayesian model with a mixture Gaussian and Laplace prior^21^. Whereas, the BSLMM^18^ used by FUSION^12^ is a combination of Bayesian variable selection model (BVSR)^22^ and linear mixed model (LMM)^23^ by assuming a normal mixture prior. However, a parametric prior is assumed for the cis-eQTL effect-sizes by both Elastic-Net and BSLMM, which restricts the capability of PrediXcan and FUSION for handling the underlying complex genetic architecture of transcriptomes. Existing studies^11; 12^ have also shown that both PrediXcan^11^ and FUSION^12^ estimated the average regression *R*^2^ (i.e., the percentage of gene expression variation that can be explained by cis-genotypes) as ~5% for human whole blood transcriptome, while the average genome-wide heritability of gene expression in human whole blood transcriptome is estimated to be more than double that quantity^24; 25^. As shown in the result section of this paper, similar underestimation of regression *R*^2^ by the Elastic-Net model (used by PrediXcan^11^) was observed in both simulated and real datasets.

Therefore, to flexibly model cis-eQTL distributions, we propose to use a nonparametric Bayesian method, where the prior for effect-sizes is nonparametric and can be estimated from the data by assuming a Dirichlet process prior on effect-size variance. This Bayesian model is also known as latent Dirichlet process regression (DPR) model^26^, which can flexibly model the underlying complex genetic architecture of transcriptomes. Thus, DPR is a more generalized model that includes Elastic-Net (implemented in PrediXcan^11^) and BSLMM (implemented in FUSION^12^) as special cases. Consequently, DPR can robustly estimate cis-eQTLs and then improve imputation *R*^2^ (the squared Pearson correlation between the observed and imputed values on test samples). Moreover, we will use a variational Bayesian algorithm^26–28^ as an alternative of Monte Carlo Markov Chain (MCMC)^29^ to efficiently fit the Bayesian model.

Similar to PrediXcan^11^ and FUSION^12^ methods, we can use DPR to estimate cis-eQTLs effect-sizes from a reference dataset, which can then be used for downstream TWAS using either individual-level or summary-level GWAS data. In subsequent sections, we first describe the DPR approach for estimating cis-eQTL effect-sizes from a reference dataset and how we can then use these effect-sizes for downstream TWAS using either individual-level or summary-level GWAS data. We then compare the performance of DPR with PrediXcan using both simulated data and real SNP and transcriptome data from the Religious Orders Study and Rush Memory Aging Project (ROS/MAP)^30–33^ for studying Alzheimer’s Disease (AD).

Our in-depth simulation studies demonstrated that the DPR method obtained higher imputation *R*^2^ on test samples, when the true expression heritability is <10% or when >1% variants are true causal cis-eQTL. Consequently, better imputation *R*^2^ resulted in improved power for follow-up association studies. Meanwhile, application of DPR to the ROS/MAP study imputed GReX for 57.6% more genes than PrediXcan. Using DPR, we also found a potentially associated gene *TRAPPC6A* for AD pathology indices that was missed by PrediXcan. Further, by using the transcriptomic imputation models fitted from ROS/MAP data and summary-level GWAS data generated from the International Genomics of Alzheimer’s Project (IGAP)^34^, we identified 3 known AD loci^34–38^ that potentially affect the late-onsite AD risk through transcript abundance. We conclude with a discussion of future topics and further describe our novel software tool TIGAR (**T**ranscriptome-**I**ntegrated **G**enetic **A**ssociation **R**esource) implementing the nonparametric Bayesian method for public use.

## Materials and Methods

We consider the following additive linear regression model for estimating the cis-eQTL effect-sizes from a reference study that has both genetic and transcriptomic data available,

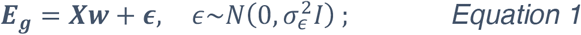

where ***E_g_*** denotes the gene expression levels (after corrections for confounding covariates such as age, sex, and principal components (PCs)) for gene *g*, ***X*** denotes the genotype matrix for all cis-genotypes (encoded as the number of minor alleles), ***w*** denotes the corresponding cis-eQTL effect-size vector, and ***ϵ*** denotes the error term. The intercept term is dropped in *Equation 1* for assuming both ***E_g_*** and ***X*** are centered at 0. Generally, SNPs within 1MB of the flanking 5’ and 3’ ends (cis-SNPs) are included in this regression model and non-zero ***ŵ*** will be used for follow-up analysis. In particular, the GReX will be imputed by

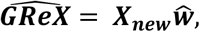

with cis-SNP data ***X_new_*** for GWAS samples.

### Nonparametric Bayesian Method

Following the latent Dirichlet process regression (DPR) model proposed in previous studies for predicting complex phenotypes^26^, we assume a normal prior 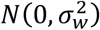 for the cis-eQTL effect-sizes (*w_i_, i* = 1,…, *p*) and a Dirichlet process (DP) prior^39^ for the effect-size variance 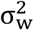 (as in *Equation 1*):

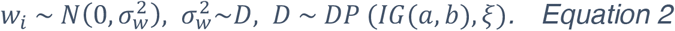

The prior distribution *D* deviates from the DP with base distribution as an Inverse Gamma (IG) distribution and concentration parameter *ξ*. Note that 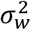 can be viewed as a latent variable and integrating out 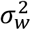 will induce a nonparametric prior distribution on *w_i_*, which is equivalent to a DP normal mixture model^26–28^,

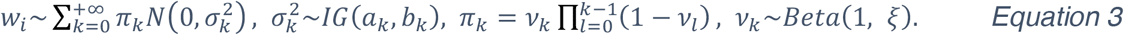

Here, the nonparametric prior distribution on *w_i_* is equivalently represented by a mixture normal prior that is a weighted sum of an infinitely number of normal distributions 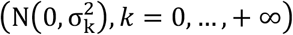, corresponding weight *π_k_* is determined by (*v_l_, l* = 0,…, *k*) with a Beta prior, and *ξ* in the Beta prior (the same concentration parameter as in Equation 2) determines the number of components with non-zero weights in the mixture normal prior. Conjugate hyper priors *ξ*~*Gamma*(*a_ξ_, b_ξ_*) and 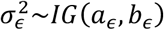 are assumed.

Generally, the hyper parameters *a_k_, b_k_, a_∊_, b_∊_* in the *Inverse Gamma* distributions can be set as 0.1 and (*a_ξ_, b_ξ_*) in the *Gamma* distribution can be set as (1, 0.1) to induce non-informative priors for 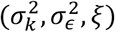. That is, the parameters 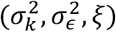 will be adaptively estimated from the data and the nonparametric prior on *w_i_* will be data-driven. The posterior estimates for ***w*** can be obtained by the MCMC^29^ or variational Bayesian algorithm^28; 40^, from the following joint conditional posterior distribution

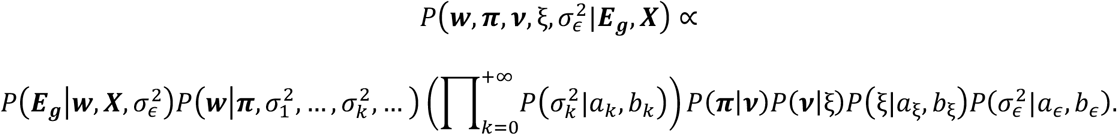

Particularly, the variational Bayesian algorithm^28; 40^ is an approximation for the MCMC^29^ with greatly improved computational efficiency, which is also used in our tool. Please refer to the supplementary note for technical details of both MCMC sampling and variational inference algorithms for obtaining the Bayesian posterior estimates for the cis-eQTL effect-sizes.

### Elastic-Net and BSLMM Methods

The Elastic-Net model^17^ (used by PrediXcan^11^) estimates the cis-eQTL effect-sizes ***ŵ*** in *Equation 1* with a combination of *L*_1_ (LASSO)^19^ and *L*_2_ (Ridge)^20^ penalties by

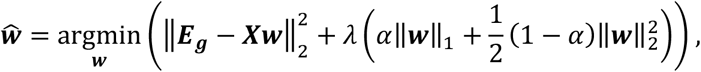

where ║·║_2_ denotes *L*_2_ norm, ║·║_1_ denotes *L*_1_ norm, *α* ∈ [0, 1] denotes the proportion of *L*_1_ penalty, and *λ* denotes the penalty parameter. Particularly, PrediXcan^11^ takes *α* = 0.5 and tunes the penalty parameter *λ* by a 5-fold cross validation.

As pointed out by previous studies^17; 21^, the Elastic-Net model is equivalent to a Bayesian model with a mixture Gaussian and Laplace (mixture normal) prior for ***w***, that is, 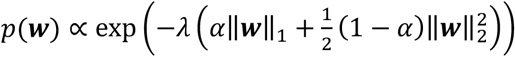. Whereas, the BSLMM^18^ assumes a mixture of two normal as the prior for cis-eQTL effect-sizes, 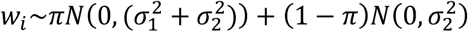. That is, the BSLMM^18^ assumes all cis-SNPs have at least a small effect, which are normally distributed with variance 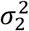, and some proportion (*π*) of cis-SNPs have an additional effect, normally distributed with variance 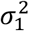. Particularly, with 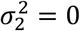, BSLMM becomes BVSR^22^, and with *π* = 0, the BSLMM becomes the LMM^23^. Therefore, the DP normal mixture^26–28^ as assumed by the DPR method includes the parametric (mixture normal) priors for Bayesian Elastic-Net^21^ and BSLMM^18^ as special cases, which is the main reason why DPR is a more generalized model including Elastic-Net and BSLMM as special cases. Thus, we believe the DPR method can robustly model complex genetic architecture and then improve the imputation *R*^2^.

### Association Study with Univariate Phenotype

Given individual-level GWAS data (genotype data ***X**_new_*, phenotype ***Y***, and covariant matrix ***C***) and cis-eQTL effect-size estimates ***ŵ***, the follow-up TWAS (using a burden type gene-based test^41^) is to test for association between 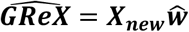 and ***Y*** based on the following generalized linear regression model

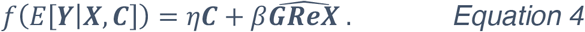

Here, *f*(·) is a pre-specified link function, which can be set as identify function for quantitative phenotype or set as logit function for dichotomous phenotype. The gene-based association test is equivalent to test *H*_0_: *β* = 0 in Equation 4.

If only summary-level data are available, we take the same approach as implemented by the FUSION^12^ method. Let ***Z*** denote the vector of Z-scores by single variant tests (Wald, likelihood ratio, score tests, etc.) for all cis-SNPs. The burden Z-score for gene-based association test is defined as

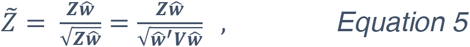

where ***V*** denotes the covariance matrix across study SNPs that can be estimated from training data or reference panels such as 1000 Genome Project^42^.

### Association Study with Multivariate Phenotype

To test for association between multivariate phenotypes and imputed GReX of the focal gene, we take a similar approach as the MultiPhen method^43^ using individual-level data. For example, consider two phenotypes (***Y***_1_, ***Y***_2_) and a covariate matrix ***C***, we first adjust for the covariates by taking the residuals (***Ỹ***_1_, ***Ỹ***_2_) respectively from the linear regression models ***Y_j_*** = ***ηC*** + ***∊***, *j* = 1,2. Then we test if the regression *R*^2^ is significantly greater than zero (*H*_0_: *R*^2^ = 0) for the following regression model

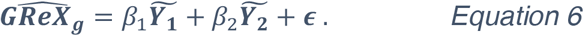

That is, we test if the multivariate phenotypes can jointly explain a non-zero percentage of variance in the imputed GReX. The p-value can be calculated by using the F-statistic for the regression *R*^2^ in *Equation 6*.

Even when only summary-level GWAS data are available, we can first obtain a burden Z-score per phenotype from *Equation 5*, i.e., 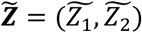 with two phenotypes. Then a similar burden approach can be used to obtain a joint Z-score for multi-phenotype test,

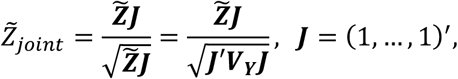

where ***V_Y_*** is the covariance matrix among study traits.

### Simulation Study Design

We conducted in-depth simulation studies to compare the performance of both PrediXcan and DPR methods with respect to imputation *R*^2^ in the test data and the power of association studies. Specifically, we used 499 ROS/MAP participants^44^ with both RNA-sequencing and genotype data as training data, and genotype data from an additional 1,200 ROS/MAP participants^44^ as test data. The test sample size (1,200) was chosen arbitrarily (randomly selected from the ROS/MAP study) to be comparable with the sample size (1,164) in the real association study of AD pathology indices. The genotyped and imputed genetic data for 2,799 cis-SNPs (with minor allele frequency (MAF) > 5% and Hardy-Weinberg p-value > 10^−5^) of the arbitrarily chosen gene *ABCA7* (see Figure S1 for the LD block structure) were used to simulate gene expression levels.

We performed comprehensive scenarios that varied the proportion of causal SNPs (out of 2,799 SNPs, influenced gene expression) among values in the vector *p_causal_* = (0.01,0.05,0.1,0.2). We varied the proportion of gene expression variance explained by causal SNPs (i.e., expression heritability), along with the proportion of phenotypic variance explained by simulated gene expression levels (i.e., phenotypic heritability), among values in the vector 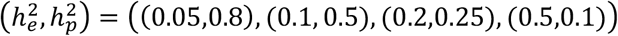 The phenotypic heritability was selected arbitrarily with respect to expression heritability such that the follow-up association study power fell within the range of (25%, 85%). We also considered various training sample sizes (100, 300, 499) for simulation scenario with *p_causal_* = 0.2 and 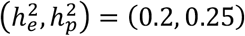.

With genotype matrix ***X_g_*** of the randomly selected causal SNPs (according to *p_causal_*), we generated effect-sizes *w_i_* from *N*(0, 1) and then re-scaled the effect-sizes to ensure the targeted 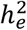. Gene expression levels were generated by ***E_g_*** = ***X_g_w*** + ∊, with 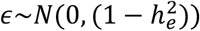. Then the phenotype values were generated by ***Y*** = *β**E_g_*** + *∊*, where *β* was selected with respect to 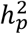 and 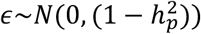.

For each scenario, we repeated simulations for 1,000 times, where we applied both PrediXcan^11^ and DPR methods to obtain imputation models with training samples, impute the GReX for test samples, and then conduct follow-up association studies using the imputed GReX. We did not compare with FUSION^12^ using BSLMM because of the computational burden of estimating cis-eQTL effect-sizes by MCMC (~2 hours per gene). The association study power was calculated as the proportion of 1,000 repeated simulations with p-value < 2.5 × 10^−6^ (genome-wide significance threshold adjusting for testing 20K independent genes).

### ROS/MAP Data

Samples in the ROS/MAP data were collected from participants of the Religious Orders Study (ROS) and the Rush Memory and Aging Project (MAP), which are prospective cohort studies of studying aging and dementia^30; 31; 33^. The ROS/MAP study recruited senior adults without known dementia at enrollment, who underwent annual clinical evaluation. Brain autopsy was done at the time of death for each participant. All participants signed an informed consent and Anatomic Gift Act, and the studies were approved by the Institutional Review Board of Rush University Medical Center, Chicago, IL. Specifically, microarray genotype data generated for 2,093 European-decent participants^44^, were further imputed to the 1000 Genome Project Phase 3^42^ in our analysis. The post-mortem brain samples (gray matter of the dorsolateral prefrontal cortex) from ~30% these participants were profiled for transcriptomic data by next-generation RNA-seqencing^45^. In this paper, we conducted gene-based association studies for two important indices of AD pathology that were quantified with *β*-antibody specific immunostains^30; 31; 33^: neurofibrillary tangle density (tangles) with stereology and *β*-amyloid load (amyloid) with image analysis. The neurofibrillary tangle density quantifies the average Tau tangle density within two or more 20μm sections from eight brain regions — hippocampus, entorhinal cortex, midfrontal cortex, inferior temporal, angular gyrus, calcarine cortex, anterior cingulate cortex, and superior frontal cortex. The *β*-amyloid load quantifies the average percent area of cortex occupied by *β*-amyloid protein in adjacent sections from the same eight brain regions.

## Results

### Simulation Studies

In the simulation studies, we observed that the DPR method performed robustly with respect to different causal proportions and gene expression heritability. Especially, when *P_causal_* > 0.01 DPR outperformed PrediXcan across all expression heritability values, giving higher imputation *R*^2^ in test data (Figure 1A). For example, when *p_causal_* = 0.2, the average imputation *R*^2^ of 1,000 simulations was estimated as 4.54% by using DPR versus 2.64% by using PrediXcan with 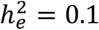, while the average imputation *R*^2^ was estimated as 12.02% by using DPR versus 9.13% by using PrediXcan with 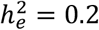 (Table 1). When *p_causal_* = 0.01, DPR performed similarly as PrediXcan with 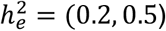 but slightly out-performed PrediXcan with 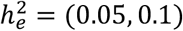 (Table 1, Figure 1).

**Figure 1.**
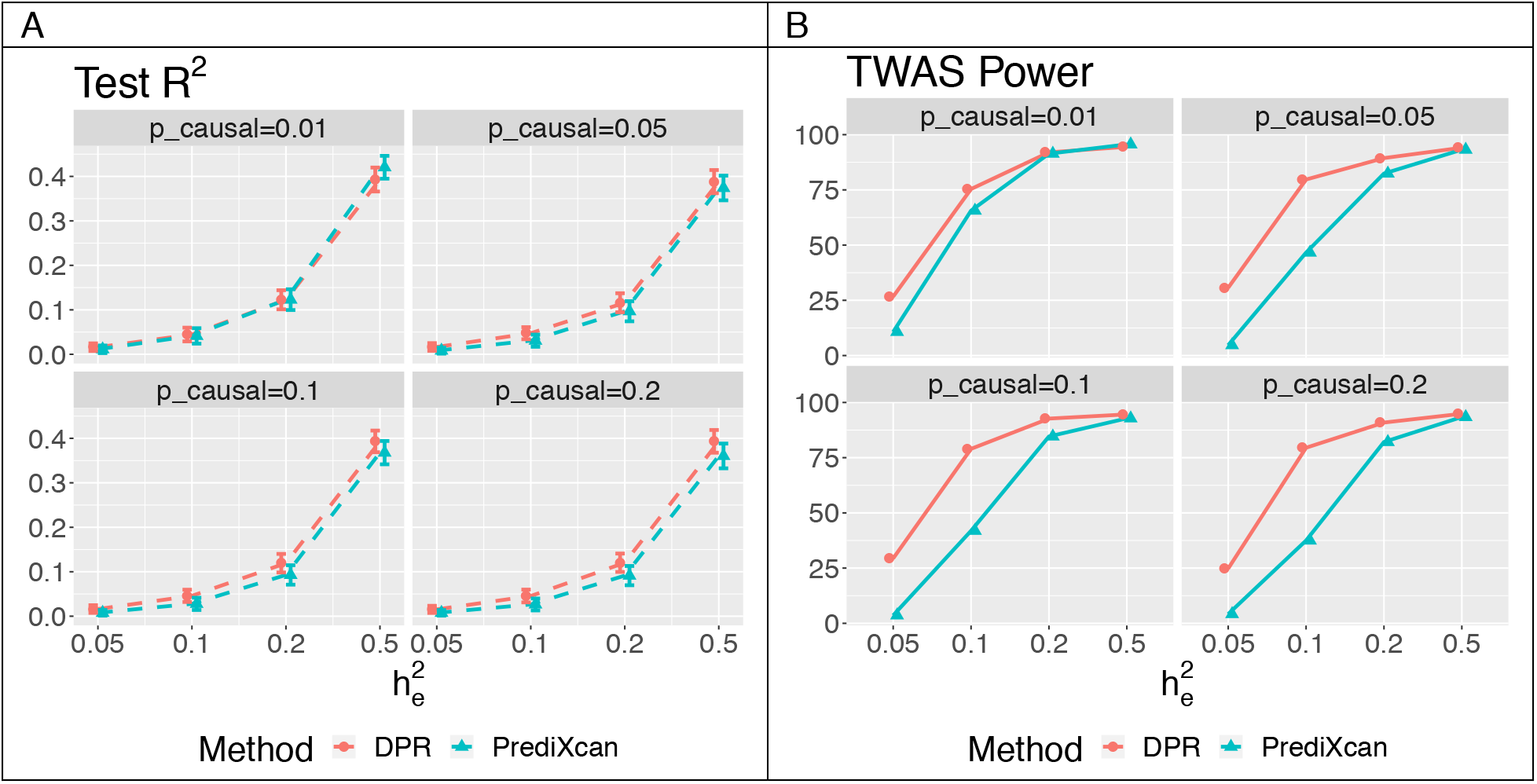
Plots of average imputation R^2^ (**A**) and TWAS power (**B**) in test samples by DPR and PrediXcan, with various proportions of true causal SNPs p_causal_ = (0.01, 0.05, 0.1, 0.2) and true expression heritability 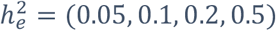. TWAS Power was evaluated with paired expression and phenotype heritability 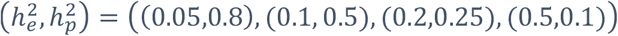.

**Table 1.** Average imputation R^2^ in the test data under various simulation scenarios, when the proportion of true causal SNPs p_causal_ = (0.01, 0.2) and expression heritability 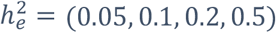. The best imputation R^2^ in the test samples are bold.

| $h_e^2$ | Causal Proportion 0.01 | | Causal Proportion 0.2 | |
| --- | --- | --- | --- | --- |
|  | DPR | PrediXcan | DPR | PrediXcan |
| 0.05 | **1.60%** | 1.12% | **1.54%** | 0.76% |
| 0.1 | **4.44%** | 4.13% | **4.54%** | 2.64% |
| 0.2 | 12.26% | **12.29%** | **12.02%** | 9.13% |
| 0.5 | 39.30% | **42.05%** | **39.32%** | 36.04% |

Consequently, when *p_causal_* > 0.01 and 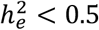, or *p_causal_* = 0.01 and 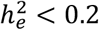, the power of association studies was higher by using DPR versus using PrediXcan imputation models (Figure 1B). When 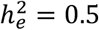, using both imputation models led to comparable power for association studies (Figure 1B). Even though both methods had similar over-estimated training *R*^2^ (Figure S2), the DPR method still resulted in higher imputation *R*^2^ for test data (Table 1; Figures 1A) and higher power for association studies (Figure 1B). In addition, from the simulation studies with various training sample sizes (100, 300, 499) and *p_causal_* = 0.2 and 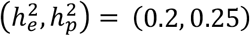, the imputation *R*^2^ and TWAS power increases as sample size increases while the DPR method consistently outperforms PrediXcan (Figure 2). Overall, these results demonstrated the advantages of the DPR method for modeling the complex genetic architecture of transcriptomes, especially when the causal proportions > 0.01 or the expression heritability < 0.2.

**Figure 2.**
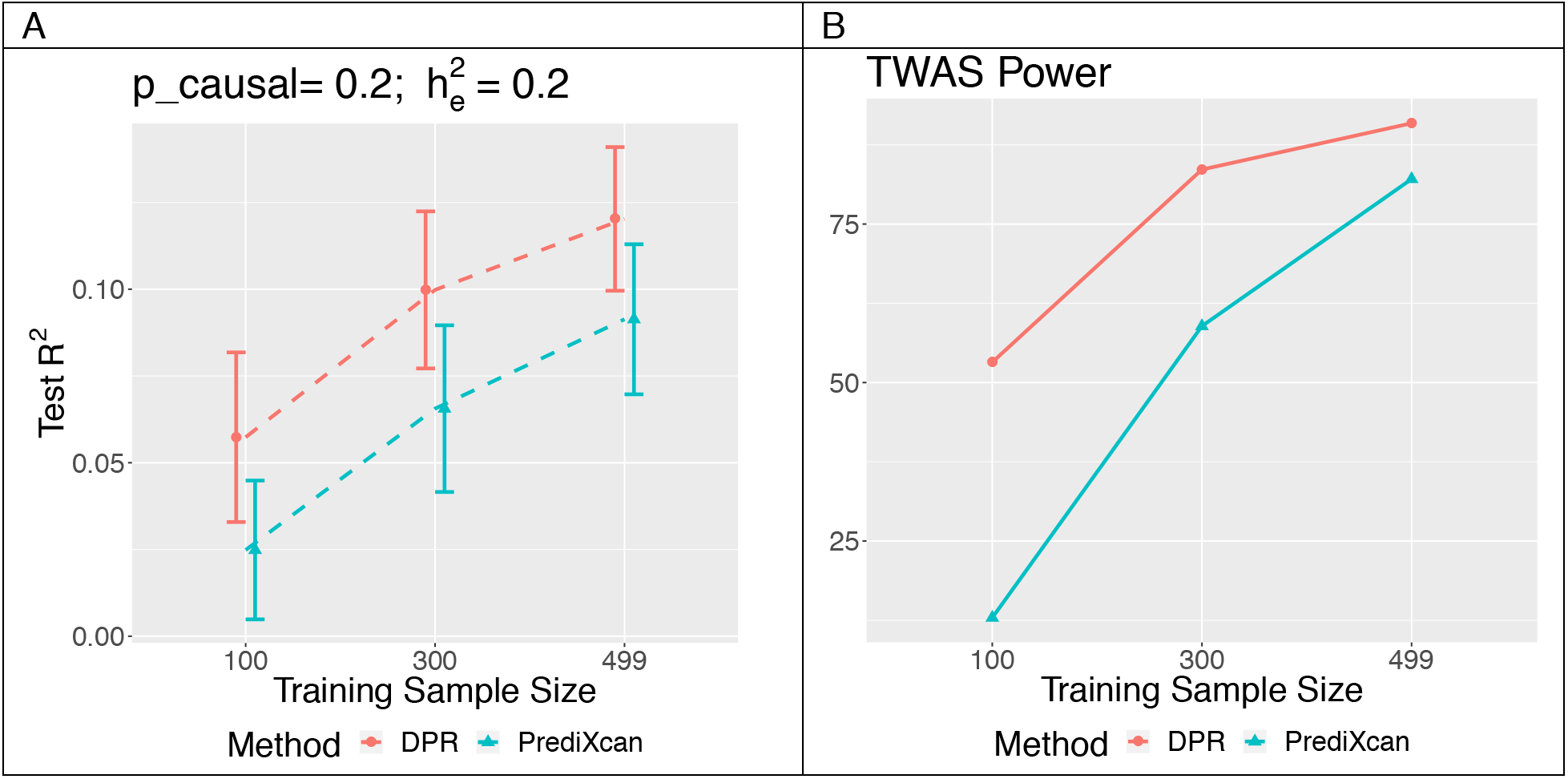
Test R^2^ (**A**) and TWAS power (**B**) from simulation studies with causal proportion p_causal_ = 0.2, expression heritability and phenotype heritability 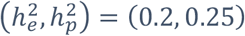, and various training sample sizes (100, 300, 499).

### Real Applications to ROS/MAP Data

To illustrate the performance of the DPR method in real studies, we applied both DPR and PrediXcan on the ROS/MAP data (see Methods). We trained the gene expression imputation models using 499 samples that have both transcriptomic data for frontal cortex tissues and genotype data (imputed to 1000 Genome Phase 3, with MAF > 5%, Hardy-Weinberg p-value > 10^−5^, and genotype imputation *R*^2^ > 0.3). A total of 15,583 genes had gene expression levels after standard RNA-sequencing quality control. The gene expression levels were first adjusted for age at death, sex, postmortem interval, study (ROS or MAP), batch effects, RNA integrity number scores, and cell type proportions (with respect to oligodendrocytes, astrocytes, microglia, neurons) by linear regression models. For each gene, cis-SNPs within the 1MB of the flanking 5’ and 3’ ends were used in the imputation models as predictors.

In this real study, we compared transcriptome-wide 5-fold cross validation (CV) regression *R*^2^ estimated by using both DPR and PrediXcan methods. Specifically, we randomly split 499 training samples into 5 folds, where the imputation *R*^2^ of each fold was calculated using the model trained with the other 4 folds samples. If the training model is null, we take the imputation *R*^2^ as 0 and take the average imputation *R*^2^ across all 5-fold test samples as 5-fold CV *R*^2^. The transcriptome-wide median of 5-fold CV *R*^2^ is 0.013 by DPR versus 0.005 by PrediXcan. The 5-fold CV *R*^2^ was used as the criterion for selecting significant imputation models (*R*^2^ > 0.01 as used by previous studies^11; 46^). From Figure 3A, we can see that the DPR method obtained more imputation models and higher imputation *R*^2^ when 5-fold CV *R*^2^ is in the range of (0.01, 0.05), which is also consistent with our simulation studies. Overall, the DPR method obtained significant imputation models for 8,742 genes versus 5,547 genes by PrediXcan (with 57.6% increases). Thus, the DPR method featuring data-driven nonparametric prior for the cis-eQTL is preferred in real studies for identifying more genes with imputable expression levels.

**Figure 3.**
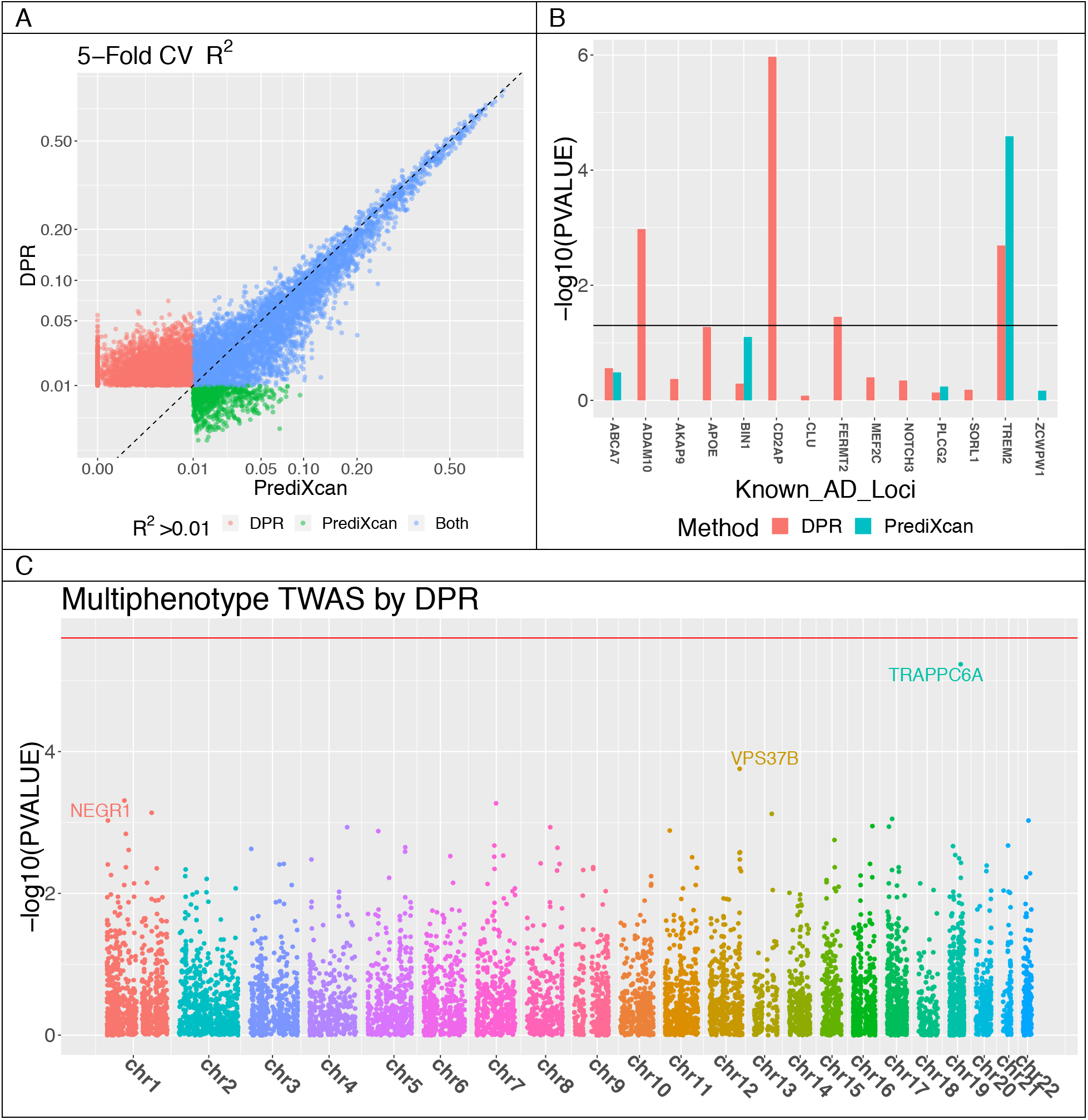
Transcriptome-wide 5-fold cross validation R^2^ (**A**) by PrediXcan and DPR with 499 ROS/MAP training samples, with different colors denoting whether the imputation R^2^ > 0.01 by DPR, PrediXcan, or both methods (genes with R^2^ < 0.01 by both DPR and PrediXcan were excluded from the plot). TWAS results (**B**) at known AD loci using GWAS summary-level statistics from IGAP and imputation models fitted from ROS/MAP data, where missing values are due to NULL imputation models by PrediXcan. Manhattan plot (**C**) for the multiphenotype TWAS (with neurofibrillary tangle density and β-amyloid load), using individual-level ROS/MAP data.

Next, we used all 499 training samples to fit imputation models for genes with respective 5-fold CV *R*^2^ > 0.01 by both DPR and PrediXcan, and then used these models to impute the GReX for all GWAS samples. We conducted univariate phenotype association studies (Methods) using all GWAS samples (N = 1,164) that have the AD pathology indices (neurofibrillary tangle density and *β*-amyloid load, with Pearson correlation 0.48) quantified. Possible confounding covariates including age at death, sex, study (ROS or MAP), smoking, education, and first 3 genotype principle components were adjusted in the association studies. Interestingly, the association studies for both AD pathology indices using the DPR imputation models identified the same top significant gene *TRAPPC6A* (within the 2MB region from the major risk gene *APOE*, encoding apolipoprotein E, but independent of *APOE*) with p-values 1.64 × 10^−5^ and 5.35 × 10^−5^ (Figure S3A and Figure S4A). Moreover, the multivariate phenotype association studies (Methods) for both AD pathology indices also identified *TRAPPC6A* as the most significant gene with p-value 5.81 × 10^−6^ and FDR 0.08 (Figure 3C). On the other hand, the PrediXcan failed to obtain a transcriptomic imputation model for *TRAPPC6A* (Figure S3B, Figure S4B, and Figure S6). Quantile-Quantile plots for these TWAS p-values were presented in Figure S5.

In addition, for 14 known common and rare loci of late-onset AD^34–38^ with significant imputation models, we conducted association studies using transcriptomic imputation models (DPR and PrediXcan) fitted from ROS/MAP data and standard GWAS results from IGAP^34^. Using the imputation models fit by DPR, we identified 3 significant loci with FDR < 0.05 (Figure 3B) — *ADAM10, CD2AP*, and *TREM2* — that potentially affect late-onsite AD risk through transcript abundance. Here, *TREM2* was also identified by using the PrediXcan imputation model (Figure 3B). Particularly, the PrediXcan method only imputed GReX for 5 out of these 14 loci. In summary, these results show that the DPR method has superior power for follow-up TWAS.

## Discussion

In this paper, we propose an improved nonparametric Bayesian method for transcriptomic data imputation, accompanied with an integrated tool (freely available on GITHUB) that is referred as **T**ranscriptome-**I**ntegrated **G**enetic **A**ssociation **R**esource (**TIGAR**). TIGAR integrates both nonparametric Bayesian (DPR) and Elastic-Net models as two options for transcriptomic data imputation, along with TWAS options using individual-level and summary-level GWAS data for univariate and multi-variate phenotypes. We show the usefulness of the DPR method by both in-depth simulations and real applications using individual-level ROS/MAP^30–33^ and summary-level IGAP^34^ data. We demonstrate that DPR is preferred over PrediXcan because of the flexible nonparametric modeling of cis-eQTL effect-sizes that results in improved imputation *R*^2^ for gene expression levels and higher power for TWAS.

With respect to user-friendly interface and computational efficiency, TIGAR can: (i) Take standard input files such as genotype files in VCF and dosage formats, phenotype files in PED format, and a combined file (tab delimited) for gene annotations and expression levels; (ii) Load input data per gene by TABIX for memory efficiency; (iii) Filter SNPs by given MAF and Hardy-Weinberg p-value thresholds; (iv) Provide options of training both Elastic-Net (using Python scripts) and DPR (using the executable tool developed with C++^26^) imputation models with unified output format; (v) Implement multi-threaded computation to take full advantage of multi-core clusters. These features make TIGAR a preferred tool for saving tedious data preparation and computation time. For example, TIGAR can complete training imputation models for ~20K genes and ~1K samples within ~20 hours and TWAS within ~1hour with a 2.4GHz 16-core CPU. The computation time is expected to increase linearly with respect to the sample size.

It is important to notice that imputing GReX with cis-eQTL effect-sizes estimated from a training dataset is analogous to the idea of estimating polygenic risk scores (PRS)^47^. Even though studies of population heterogeneity are lacked for imputing GReX, the same philosophy of estimating PRS still applies because of the same underlying statistical models. That is, given both genetic and transcriptomic heterogeneities across different populations, one need to be cautious not using training dataset of a different ethnicity for TWAS^47^.

As observed in the real ROS/MAP studies, there remains a large gap between the 5-fold CV *R*^2^ using cis-eQTL predictors (~5%) and the average genome-wide heritability of gene expression levels (21.8% estimated by GCTA^48^ based on a LMM). This is likely due to the large trans-acting contribution to transcript abundance documented for most genes. Thus, we hypothesize that it is promising to further improve the imputation *R*^2^ by fitting transcriptomic imputation models with genome-wide variants as predictors. Scalable Bayesian inference techniques such as the Expectation Maximization MCMC (EM-MCMC) algorithm^49^ are required for incorporating genome-wide variants.

Another limitation of existing TWAS methods is that the uncertainty of cis-eQTL effect-size estimates has not been taken into accounted in the association studies. A Bayesian framework can also be derived by taking the standard errors of these cis-eQTL effect-size estimates as prior standard deviations, which is part of our continuing research.

Besides the follow-up gene-based association studies (i.e., TWAS) described in this paper, the transcriptomic imputation models can be further extended by incorporating environmental contributions. The imputed transcript abundance levels can then be used for gene network analysis, differential gene expression analysis, and transcriptome mediation analysis with GWAS data. Validation of transcriptomic prediction accuracy in independent datasets will be critical in this regard, but unfortunately multiple large and similar datasets are not yet generally available for tissues other than peripheral blood.

In conclusion, we expect our work will provide a convenient and improved tool for transcriptomic imputation using the currently available rich reference datasets, as well as enhanced gene mapping for better understanding the genetic etiology of complex traits.

## Supplemental Data

Supplementary Text and Figures

## Acknowledgments

J.Y. was supported by the startup funding from Emory University Department of Human Genetics. A.P.W. and T.S.W. were supported by National Institutes of Health (NIH) R01AG056533. M.P.E. was supported by NIH R01GM11796. L.C.T. was supported by the Dermatology Foundation, the Arthritis National Research Foundation, the National Psoriasis Foundation, and NIH K01AR072129. ROS/MAP study data were provided by the Rush Alzheimer’s Disease Center, Rush University Medical Center, Chicago, IL. Data collection was supported through funding by NIA grants P30AG10161, R01AG15819, R01AG17917, R01AG30146, R01AG36836, U01AG32984, U01AG46152, the Illinois Department of Public Health, and the Translational Genomics Research Institute. In addition, we thank Thanneer Perumal and Benjamin Logsdon for performing quality control of the ROS/MAP RNA-sequencing data and for creating the brain cell type proportions.

## Web Resources

TIGAR, https://github.com/yanglab-emory/TIGAR

PrediXcan, https://github.com/hakyim/PrediXcan

FUSION, http://gusevlab.org/projects/fusion/

ROS/MAP data, https://www.synapse.org/#!Synapse:syn3219045; http://www.radc.rush.edu/

IGAP data, http://web.pasteur-lille.fr/en/recherche/u744/igap/igap_download.php

