## Supplementary text and figures for "TIGAR: An Improved Bayesian Tool for Transcriptomic Data Imputation Enhances Gene Mapping of Complex Traits"

### 1 Supplemental Text: Computational Algorithms for the Nonparametric Bayesian Method

#### 1.1 Nonparametric Bayesian Dirichlet Process Regression Model

We consider the following additive linear regression model for estimating the cis-eQTL effect-sizes,

$$\mathbf{E_g} = \mathbf{X_{n\times p}w_{p\times 1}} + \boldsymbol{\varepsilon},\ \boldsymbol{\varepsilon} \sim \mathbf{N}(\mathbf{0}, \sigma_\epsilon^{\mathbf{2}}\mathbf{I}),\ \sigma_\epsilon^{\mathbf{2}} \sim \mathbf{IG}(\mathbf{a}_\epsilon, \mathbf{b}_\epsilon), \tag{1}$$

where vector $\mathbf{E_g}$ denotes the gene expression levels for gene $g$, $\boldsymbol{X_{n\times p}}$ denotes the genotype matrix for $n$ samples and $p$ cis-SNPs (encoded as the number of minor alleles), $\boldsymbol{w}$ denotes the corresponding cis-eQTL effect-size vector, vector $\varepsilon$ denotes the error term, and $\boldsymbol{I}$ denotes an identity matrix. The intercept term is dropped in model (1) for assuming both $\mathbf{E_g}$ and $\boldsymbol{X}$ are centered at 0. The error variance $\sigma_\epsilon^2$ is assumed with an Inverse Gamma (IG) prior distribution.

Following the latent Dirichlet process regression (DPR) model [1], we assume a normal prior $N(0, \sigma_\epsilon^2\sigma_w^2)$ for the cis-eQTL effect-sizes and a Dirichlet process (DP) prior [2] for $\sigma_w^2$. That is,

$$w_i \sim N(0, \sigma_\epsilon^2\sigma_w^2),\ \sigma_w^2 \sim D,\ D \sim DP(IG(a,b), \xi),\ i = 1, \cdots, p, \tag{2}$$

where the prior distribution $D$ deviates from the $DP$ with base distribution as an Inverse Gamma distribution and concentration parameter $\xi$. In particular, different from the notation in the main text, the prior effect-size variance is assumed to be scaled by the inverse of the error variance $(\sigma_\epsilon^2)^{-1}$ for computational simplicity. Here, $\sigma_w^2$ can be viewed as a latent variable and integrating out $\sigma_w^2$ will induce a nonparametric prior distribution on $w_i$, which is equivalent to the following DP normal mixture model [3, 4],

$$\begin{aligned} w_i &\sim \pi_0 N(0, \sigma_\epsilon^2\sigma_0^2) + \sum_{k=1}^{+\infty}\pi_k N(0, \sigma_\epsilon^2(\sigma_k^2 + \sigma_0^2)),\ \pi_k = \nu_k\prod_{l=0}^{k-1}(1-\nu_l); \\ \nu_k &\sim Beta(1, \xi),\ \xi \sim Gamma(a_\xi, b_\xi),\ \sigma_k^2 \sim IG(a_k, b_k),\ k = 0, 1, \cdots, +\infty. \end{aligned} \tag{3}$$

Here, the nonparametric prior distribution on $w_i$ is equivalently represented by a mixture normal prior that is a weighted sum of an infinitely number of zero-mean normal distributions. The variance terms of these normal distributions are assumed to be scaled by the inverse of the error variance $(\sigma_\epsilon^2)^{-1}$ and centered by $-\sigma_0^2$ for those corresponding to $k > 0$. Note that the $\sigma_0^2$ can also be viewed as the smallest variance component, such that other variance components can be written as $(\sigma_k^2 + \sigma_0^2)$. The weight parameter $\pi_k$ is determined by $\{\nu_l, l = 0, \cdots, k\}$ with a Beta prior, where the parameter $\xi$ determines the number of components with non-zero weights ($\xi$ is the same concentration parameter as in (2)). Generally, the hyper parameters $\{a_k, b_k, a_\epsilon, b_\epsilon\}$ in the Inverse Gamma distributions can be set as 0.1 and $(a_\xi, b_\xi)$ in the Gamma distribution can be set as $(1, 0.1)$ to induce non-informative priors for $(\sigma_k^2, \sigma_\epsilon^2, \xi)$.

For computational convenience, following previous Bayesian models [1, 5], we group the effect-sizes corresponding to the first normal component that has the smallest variance term, $N(0, \sigma_\epsilon^2 \sigma_0^2)$, as a random effect term $\boldsymbol{u}$. Then the above model (1, 3) is equivalent to

$$\begin{aligned} \mathbf{E_g} &= \mathbf{X}\widetilde{\mathbf{w}} + \mathbf{u} + \boldsymbol{\varepsilon};\ \mathbf{u} \sim \mathbf{N}(\mathbf{0}, \sigma_\epsilon^{\mathbf{2}}\sigma_{\mathbf{0}}^{\mathbf{2}}\mathbf{K}),\ \boldsymbol{\varepsilon} \sim \mathbf{N}(\mathbf{0}, \sigma_\epsilon^{\mathbf{2}}\mathbf{I}),\ \sigma_\epsilon^{\mathbf{2}} \sim \mathbf{IG}(\mathbf{a}_\epsilon, \mathbf{b}_\epsilon); \quad (4) \\ \widetilde{w_i} &= \sum_{k=1}^{+\infty} \pi_k N(0, \sigma_\epsilon^2 \sigma_k^2),\ \pi_k = \nu_k \prod_{l=0}^{k-1}(1-\nu_l),\ k = 1, \cdots, +\infty; \\ \nu_k &\sim Beta(1, \xi),\ \xi \sim Gamma(a_\xi, b_\xi);\ \sigma_k^2 \sim IG(a_k, b_k);\ k = 0, 1, , \cdots, +\infty; \end{aligned}$$

where $\boldsymbol{K} = \boldsymbol{X}\boldsymbol{X'}/p$ is the Genetic Relatedness Matrix (GRM) [1, 5]. Note that the random effect term can be written as

$$\boldsymbol{u} = \boldsymbol{X}\boldsymbol{\zeta};\ \zeta_i \sim N(0, \sigma_\epsilon^2 \sigma_0^2/p),\ i = 1, ..., p; \quad (5)$$

where $\zeta_i$ denotes the random effect-size for SNP $i$. Our computational algorithms will be derived with respect to model (4) for estimating $(\widetilde{\boldsymbol{w}}, \boldsymbol{\zeta})$, which will then give estimates for the cis-eQTL effect-sizes $\boldsymbol{w} = \widetilde{\boldsymbol{w}} + \boldsymbol{\zeta}$ as in model (1).

Based on model (4), the joint conditional posterior function for all parameters in the model is given by

$$\begin{gathered} P(\widetilde{\boldsymbol{w}}, \boldsymbol{u}, \boldsymbol{\nu}, \xi, \sigma_\epsilon^2, \sigma_0^2, \sigma_1^2, \cdots, \sigma_k^2, \cdots | \boldsymbol{E_g}, \boldsymbol{X}, \boldsymbol{K}) \propto \quad (6) \\ P(\boldsymbol{E_g}|\boldsymbol{X}, \boldsymbol{K}, \widetilde{\boldsymbol{w}}, \boldsymbol{u}, \sigma_\epsilon^2) P(\boldsymbol{w}|\boldsymbol{\nu}, \sigma_1^2, \cdots, \sigma_k^2, \cdots) P(\boldsymbol{u}|\sigma_\epsilon^2, \sigma_0^2, \boldsymbol{K}) \\ \left(\prod_{k=0}^{+\infty} P(\sigma_k^2|a_k, b_k)\right) P(\boldsymbol{\nu}|\xi) P(\xi|a_\xi, b_\xi) P(\sigma_\epsilon^2|a_\epsilon, b_\epsilon). \end{gathered}$$

One convenience for considering model (4) is that one can integrate out the random effect term $\boldsymbol{u}$ from (6) and then implement Gibbs sampling [6] for improved mixing in the Markov Chain Monte Carlo (MCMC) sampling. Specifically,

$$\begin{gathered} P(\widetilde{\boldsymbol{w}}, \boldsymbol{\nu}, \xi, \sigma_\epsilon^2, \sigma_0^2, \sigma_1^2, \cdots, \sigma_k^2, \cdots | \boldsymbol{E_g}, \boldsymbol{X}, \boldsymbol{K}) = \int P(\widetilde{\boldsymbol{w}}, \boldsymbol{u}, \boldsymbol{\nu}, \xi, \sigma_\epsilon^2, \sigma_0^2, \sigma_1^2, \cdots, \sigma_k^2, \cdots | \boldsymbol{E_g}, \boldsymbol{X}, \boldsymbol{K}) d\boldsymbol{u} \\ = \int P(\boldsymbol{E_g}|\boldsymbol{X}, \boldsymbol{K}, \widetilde{\boldsymbol{w}}, \boldsymbol{u}, \sigma_\epsilon^2) P(\boldsymbol{u}|\sigma_\epsilon^2, \sigma_0^2, \boldsymbol{K}) d\boldsymbol{u} \times \\ P(\boldsymbol{w}|\boldsymbol{\nu}, \sigma_1^2, \cdots, \sigma_k^2, \cdots) \left(\prod_{k=0}^{+\infty} P(\sigma_k^2|a_k, b_k)\right) P(\boldsymbol{\nu}|\xi) P(\xi|a_\xi, b_\xi) P(\sigma_\epsilon^2|a_\epsilon, b_\epsilon), \quad (7) \end{gathered}$$

where

$$\begin{aligned} \int P(\boldsymbol{E_g}|\boldsymbol{X}, \boldsymbol{K}, \widetilde{\boldsymbol{w}}, \boldsymbol{u}, \sigma_\epsilon^2) P(\boldsymbol{u}|\sigma_\epsilon^2, \sigma_0^2, \boldsymbol{K}) du &\propto |\sigma_\epsilon^2 \boldsymbol{H}|^{-\frac{1}{2}} \exp\left\{-\frac{1}{2\sigma_\epsilon^2}(\boldsymbol{E_g} - \boldsymbol{X}\widetilde{\boldsymbol{w}})' \boldsymbol{H}^{-1}(\boldsymbol{E_g} - \boldsymbol{X}\widetilde{\boldsymbol{w}})\right\}, \\ \boldsymbol{H} &= \boldsymbol{I} + \sigma_0^2 \boldsymbol{K}. \end{aligned}$$

The following MCMC Sampling and Variational Inference algorithms will be derived based on the joint conditional posterior density function (7).

##### 1.2 MCMC Sampling

To facilitate MCMC sampling, for each SNP $i$, we assume a corresponding indicator vector $\boldsymbol{\gamma_i} = \{\gamma_{ik} \in \{0,1\}, k = 1, ..., +\infty\}$ that indicates if the $k$th normal component contributes to the effect-size distribution. Consequently, $\gamma_{ik}$ has a $Bernoulli(\pi_k)$ prior. Let $\boldsymbol{\gamma}$ denote the indicator vectors $\{\boldsymbol{\gamma_i}, i = 1, \cdots, p\}$. Consequently, the joint conditional posterior density function (7) becomes

$$\begin{gathered}
P(\widetilde{\boldsymbol{w}}, \boldsymbol{\gamma}, \boldsymbol{\nu}, \xi, \sigma_\epsilon^2, \sigma_0^2, \sigma_1^2, \cdots, \sigma_k^2, \cdots | \boldsymbol{E_g}, \boldsymbol{X}, \boldsymbol{K}) \propto \\
|\sigma_\epsilon^2 \boldsymbol{H}|^{-\frac{1}{2}} \exp\left\{-\frac{1}{2\sigma_\epsilon^2}(\boldsymbol{E_g} - \boldsymbol{X}\widetilde{\boldsymbol{w}})'\boldsymbol{H}^{-1}(\boldsymbol{E_g} - \boldsymbol{X}\widetilde{\boldsymbol{w}})\right\} \times \\
P(\boldsymbol{w}|\boldsymbol{\gamma}, \sigma_\epsilon^2, \sigma_1^2, \cdots, \sigma_k^2, \cdots) \left(\prod_{k=0}^{+\infty} P(\sigma_k^2 | a_k, b_k)\right) P(\boldsymbol{\gamma}|\boldsymbol{\nu}) P(\boldsymbol{\nu}|\xi) P(\xi | a_\xi, b_\xi) P(\sigma_\epsilon^2 | a_\epsilon, b_\epsilon) \propto \\
|\sigma_\epsilon^2 \boldsymbol{H}|^{-\frac{1}{2}} \exp\left\{-\frac{1}{2\sigma_\epsilon^2}(\boldsymbol{E_g} - \boldsymbol{X}\widetilde{\boldsymbol{w}})'\boldsymbol{H}^{-1}(\boldsymbol{E_g} - \boldsymbol{X}\widetilde{\boldsymbol{w}})\right\} \times \\
\left(\prod_{i=1}^{p} \prod_{k=1}^{+\infty} \gamma_{ik} N(\widetilde{w_{ik}}; 0, \sigma_\epsilon^2 \sigma_k^2)\right) \left(\prod_{k=0}^{+\infty} IG(\sigma_k^2; a_k, b_k)\right) \times \\
\left(\prod_{i=1}^{p} \prod_{k=1}^{+\infty} Bernoulli(\gamma_{ik}; \pi_k = \nu_k \prod_{l=0}^{k-1}(1 - \nu_l))\right) \left(\prod_{k=0}^{+\infty} Beta(\nu_k; 1, \xi)\right) \times \\
Gamma(\xi; a_\xi, b_\xi) IG(\sigma_\epsilon^2; a_\epsilon, b_\epsilon). \qquad (8)
\end{gathered}$$

Denoting $\widetilde{w_i} = \sum_{k=1}^{+\infty} \gamma_{ik} \widetilde{w_{ik}}$, $\widetilde{w_{ik}} \sim N(0, \sigma_\epsilon^2 \sigma_k^2)$, Then the log joint conditional posterior density function is given by

$$\begin{gathered}
log(P(\widetilde{\boldsymbol{w}}, \boldsymbol{\gamma}, \boldsymbol{\nu}, \xi, \sigma_\epsilon^2, \sigma_0^2, \sigma_1^2, \cdots, \sigma_k^2, \cdots | \boldsymbol{E_g}, \boldsymbol{X}, \boldsymbol{K})) = \\
C - \frac{1}{2} log |\sigma_\epsilon^2 \boldsymbol{H}| - \frac{1}{2\sigma_\epsilon^2}(\boldsymbol{E_g} - \boldsymbol{X}\widetilde{\boldsymbol{w}})'\boldsymbol{H}^{-1}(\boldsymbol{E_g} - \boldsymbol{X}\widetilde{\boldsymbol{w}}) + \\
\sum_{i=1}^{p} \sum_{k=1}^{+\infty} \gamma_{ik} \left[-\frac{1}{2} log(\sigma_\epsilon^2) - \frac{1}{2} log(\sigma_k^2) - \frac{\widetilde{w_{ik}}^2}{2\sigma_\epsilon^2 \sigma_k^2}\right] + \sum_{k=0}^{+\infty} \left[-(a_k + 1) log(\sigma_k^2) - \frac{b_k}{\sigma_k^2}\right] + \\
\sum_{i=1}^{p} \sum_{k=1}^{+\infty} \left[\gamma_{ik} \left(log(\nu_k) + \sum_{l=0}^{k-1} log(1 - \nu_l)\right) + (1 - \gamma_{ik}) log\left(1 - \nu_k \prod_{l=0}^{k-1}(1 - \nu_l)\right)\right] + \\
\sum_{k=0}^{+\infty} [(\xi - 1) log(1 - \nu_k) + log(\xi)] + (a_\xi - 1) log(\xi) - b_\xi \xi - (a_\epsilon + 1) log(\sigma_\epsilon^2) - \frac{b_\epsilon}{\sigma_\epsilon^2}, \qquad (9)
\end{gathered}$$

where $C$ denotes a normalization constant that is free of parameters. Based on the log conditional posterior density function (9), the following MCMC sampling scheme is derived for obtaining the posterior estimates for $\widetilde{\boldsymbol{w}}$.

###### 1.2.1 Gibbs Sampling Scheme

From (9), for each parameter in the model, we derive the log conditional density function conditioning on other parameters as follows:

- $\widetilde{w_{ik}}$ **and** $\gamma_{ik}$

  The log joint conditional density function for $\widetilde{w_{ik}}, \gamma_{ik}$ is given by,

$$\begin{gathered}
log(P(\widetilde{w_{ik}}, \gamma_{ik}|\cdot)) = C - \frac{\boldsymbol{x_i H^{-1} x_i}}{2\sigma_\epsilon^2}(\widetilde{w_{ik}}^2 + 2\widetilde{w_{ik}}\widetilde{w}_{i(-k)}) + \frac{1}{\sigma_\epsilon^2}\boldsymbol{x_i'}\boldsymbol{H}^{-1}(\boldsymbol{E_g} - \sum_{j \neq i} \boldsymbol{x_j}\widetilde{w_j})\widetilde{w_{ik}} + \\
\gamma_{ik}\left[-\frac{1}{2} log(\sigma_\epsilon^2) - \frac{1}{2} log(\sigma_k^2) - \frac{\widetilde{w_{ik}}^2}{2\sigma_\epsilon^2 \sigma_k^2}\right] + \\
\gamma_{ik}\left(log(\nu_k) + \sum_{l=0}^{k-1} log(1 - \nu_l)\right) + (1 - \gamma_{ik}) log\left(1 - \nu_k \prod_{l=0}^{k-1}(1 - \nu_l)\right), \qquad (10)
\end{gathered}$$

where $\widetilde{w}_{i(-k)} = \sum_{l \neq k} \widetilde{w}_{il}$.

Then the log conditional posterior distribution for $(\widetilde{w_{ik}} | \gamma_{ik} = 1, \cdot)$ is given by

$$\begin{aligned}
log(P(\widetilde{w_{ik}}|\gamma_{ik}=1,\cdot)) &= C - \frac{\boldsymbol{x_i H^{-1} x_i}}{2\sigma_\epsilon^2}\widetilde{w_{ik}}^2 - \frac{\widetilde{w_{ik}}^2}{2\sigma_\epsilon^2\sigma_k^2} + \frac{1}{\sigma_\epsilon^2}\boldsymbol{x_i'}\boldsymbol{H}^{-1}(\boldsymbol{E_g} - \sum_{j\neq i}\boldsymbol{x_j}\widetilde{w_j} - \boldsymbol{x_i}\widetilde{w}_{i(-k)})\widetilde{w_{ik}} \\
&= C - \frac{\boldsymbol{x_i H^{-1} x_i} + \sigma_k^{-2}}{2\sigma_\epsilon^2}\widetilde{w_{ik}}^2 + \frac{1}{\sigma_\epsilon^2}\boldsymbol{x_i'}\boldsymbol{H}^{-1}(\boldsymbol{E_g} - \sum_{j\neq i}\boldsymbol{x_j}\widetilde{w_j} - \boldsymbol{x_i}\widetilde{w}_{i(-k)})\widetilde{w_{ik}}; \\
P(\widetilde{w_{ik}}|\gamma_{ik}=1,\cdot) &\sim N(m_{ik}, s_{ik}^2), \\
m_{ik} &= \frac{\boldsymbol{x_i'}\boldsymbol{H}^{-1}(\boldsymbol{E_g} - \sum_{j\neq i}\boldsymbol{x_j}\widetilde{w_j} - \boldsymbol{x_i}\widetilde{w}_{i(-k)})}{\boldsymbol{x_i H^{-1} x_i} + \sigma_k^{-2}}, \\
s_{ik}^2 &= \frac{\sigma_\epsilon^2}{\boldsymbol{x_i H^{-1} x_i} + \sigma_k^{-2}}. \qquad (11)
\end{aligned}$$

After integrating out $\widetilde{w_{ik}}$ from (10), the conditional probability for $(\gamma_{ik} = 1|\cdot)$ is given by

$$\begin{aligned}
P(\gamma_{ik}=1|\cdot) &= \pi_{ik} = \int P(\widetilde{w_{ik}}, \gamma_{ik}=1|\cdot)d\widetilde{w_{ik}} \propto \\
& \exp\left\{-\frac{1}{2}log(\sigma_\epsilon^2) - \frac{1}{2}log(\sigma_k^2) + log(\nu_k) + \sum_{l=0}^{k-1}log(1-\nu_l)\right\} \times \\
& \int \exp\left\{-\frac{\boldsymbol{x_i H^{-1} x_i} + \sigma_k^{-2}}{2\sigma_\epsilon^2}\widetilde{w_{ik}}^2 + \frac{1}{\sigma_\epsilon^2}\boldsymbol{x_i'}\boldsymbol{H}^{-1}(\boldsymbol{E_g} - \sum_{j\neq i}\boldsymbol{x_j}\widetilde{w_j} - \boldsymbol{x_i}\widetilde{w}_{i(-k)})\widetilde{w_{ik}}\right\} d\widetilde{w_{ik}} \propto \\
& \exp\left\{-\frac{1}{2}log(\sigma_\epsilon^2) - \frac{1}{2}log(\sigma_k^2) + log(\nu_k) + \sum_{l=0}^{k-1}log(1-\nu_l)\right\} \sqrt{2\pi s_k^2}\exp\{\frac{m_{ik}^2}{2s_k^2}\} \times \\
& \int \frac{1}{\sqrt{2\pi s_k^2}}\exp\left\{-\frac{1}{2s_{ik}^2}\left(\widetilde{w_{ik}}^2 - 2m_{ik}\widetilde{w_{ik}} + m_{ik}^2\right)\right\} d\widetilde{w_{ik}} \propto \\
& \sqrt{2\pi s_k^2}\exp\left\{\frac{m_{ik}^2}{2s_k^2} - \frac{1}{2}log(\sigma_\epsilon^2) - \frac{1}{2}log(\sigma_k^2) + log(\nu_k) + \sum_{l=0}^{k-1}log(1-\nu_l)\right\} \propto \\
& \exp\left\{\frac{m_{ik}^2}{2s_k^2} + log(s_k) - log(\sigma_\epsilon) - log(\sigma_k) + log(\nu_k) + \sum_{l=0}^{k-1}log(1-\nu_l)\right\}, \qquad (12)
\end{aligned}$$

where $\int \frac{1}{\sqrt{2\pi s_k^2}}\exp\left\{-\frac{1}{2s_{ik}^2}\left(\widetilde{w_{ik}}^2 - 2m_{ik}\widetilde{w_{ik}} + m_{ik}^2\right)\right\}d\widetilde{w_{ik}} = 1$ because the integrand is just the density function of $N(\widetilde{w_{ik}}; m_{ik}, s_k^2)$.

- $\nu_k$

The log conditional density function for $\nu_k$ is given by,

$$\begin{aligned}
log(P(\nu_k|\cdot)) &= \sum_{i=1}^{p}\sum_{k=1}^{+\infty}\left[\gamma_{ik}\left(log(\nu_k)+\sum_{l=0}^{k-1}log(1-\nu_l)\right)+(1-\gamma_{ik})log\left(1-\nu_k\prod_{l=0}^{k-1}(1-\nu_l)\right)\right]+ \\
&\quad \sum_{k=1}^{+\infty}(\xi-1)log(1-\nu_k)+C, \\
&= \sum_{i=1}^{p}\gamma_{ik}log(\nu_k)+\sum_{i=1}^{p}\sum_{l=k+1}^{+\infty}\gamma_{ik}log(1-\nu_k)+ \\
&\quad \sum_{i=1}^{p}\sum_{j=k}^{+\infty}(1-\gamma_{ij})log\left(1-\nu_j\prod_{l=0}^{j-1}(1-\nu_l)\right)+(\xi-1)log(1-\nu_k)+C, \\
&\approx \sum_{i=1}^{p}\gamma_{ik}log(\nu_k)+\sum_{i=1}^{p}\sum_{l=k+1}^{+\infty}\gamma_{ik}log(1-\nu_k)+(\xi-1)log(1-\nu_k)+C, \\
P(\nu_k|\cdot) &\sim Beta(\kappa_k,\xi_k),\ \kappa_k=\sum_{i=1}^{p}\gamma_{ik}+1,\ \xi_k=\sum_{i=1}^{p}\sum_{l=k+1}^{+\infty}\gamma_{il}+\xi. \qquad (13)
\end{aligned}$$

- $\sigma_k^2, k>0$
  The log conditional density function for $\sigma_k^2, k>0$ is given by,

$$\begin{aligned}
log(P(\sigma_k^2|\cdot)) &= C+\sum_{i=1}^{p}\sum_{k=1}^{+\infty}\gamma_{ik}\left[-\frac{1}{2}log(\sigma_k^2)-\frac{\widetilde{w_{ik}}^2}{2\sigma_\epsilon^2\sigma_k^2}\right]+\sum_{k=0}^{+\infty}\left[-(a_k+1)log(\sigma_k^2)-\frac{b_k}{\sigma_k^2}\right] \\
&= C+\sum_{i=1}^{p}\gamma_{ik}\left[-\frac{1}{2}log(\sigma_k^2)-\frac{\widetilde{w_{ik}}^2}{2\sigma_\epsilon^2\sigma_k^2}\right]-(a_k+1)log(\sigma_k^2)-\frac{b_k}{\sigma_k^2} \\
&= C-\left(\frac{1}{2}\sum_{i=1}^{p}\gamma_{ik}+a_k+1\right)log(\sigma_k^2)-\left(\frac{\sum_{i=1}^{p}(\gamma_{ik}\widetilde{w_{ik}}^2)}{2\sigma_\epsilon^2}+b_k\right)\frac{1}{\sigma_k^2}; \\
P(\sigma_k^2|\cdot) &\sim IG(\widetilde{a_k},\widetilde{b_k}),\ \widetilde{a_k}=\frac{1}{2}\sum_{i=1}^{p}\gamma_{ik}+a_k,\ \widetilde{b_k}=\frac{\sum_{i=1}^{p}(\gamma_{ik}\widetilde{w_{ik}}^2)}{2\sigma_\epsilon^2}+b_k. \qquad (14)
\end{aligned}$$

- $\xi$ The log conditional density function for $\xi$ is given by,

$$\begin{aligned}
log(P(\xi|\cdot)) &= C+\sum_{k=0}^{+\infty}[(\xi-1)log(1-\nu_k)+log(\xi)]+(a_\xi-1)log(\xi)-b_\xi\xi \\
&= C+\left(a_\xi+\sum_{k=0}^{+\infty}1_k-1\right)log(\xi)-\left(b_\xi-\sum_{k=0}^{+\infty}log(1-\nu_k)\right)\xi; \\
(P(\xi|\cdot)) &\sim Gamma(\widetilde{a_\xi},\widetilde{b_\xi}),\ \widetilde{a_\xi}=a_\xi+\sum_{k=0}^{+\infty}1_k,\ \widetilde{b_\xi}=b_\xi-\sum_{k=0}^{+\infty}log(1-\nu_k). \qquad (15)
\end{aligned}$$

- $\sigma_\epsilon^2$
  The log conditional density function for $\sigma_\epsilon^2$ is given by,

$$\begin{aligned}
log(P(\sigma_\epsilon^2|\cdot)) &= C - \frac{1}{2}log|\sigma_\epsilon^2 \boldsymbol{H}| - \frac{1}{2\sigma_\epsilon^2}(\boldsymbol{E_g} - \boldsymbol{X}\widetilde{\boldsymbol{w}})'\boldsymbol{H}^{-1}(\boldsymbol{E_g} - \boldsymbol{X}\widetilde{\boldsymbol{w}}) + \\
&\quad \sum_{i=1}^{p}\sum_{k=1}^{+\infty}\gamma_{ik}\left[-\frac{1}{2}log(\sigma_\epsilon^2) - \frac{\widetilde{w_{ik}}^2}{2\sigma_\epsilon^2\sigma_k^2}\right] - (a_\epsilon + 1)log(\sigma_\epsilon^2) - \frac{b_\epsilon}{\sigma_\epsilon^2} \\
&= C - log(\sigma_\epsilon^2)\left(\frac{n}{2} + \frac{1}{2}\sum_{i=1}^{p}\sum_{k=1}^{+\infty}\gamma_{ik} + a_\epsilon + 1\right) - \\
&\quad \frac{1}{\sigma_\epsilon^2}\left(\frac{1}{2}(\boldsymbol{E_g} - \boldsymbol{X}\widetilde{\boldsymbol{w}})'\boldsymbol{H}^{-1}(\boldsymbol{E_g} - \boldsymbol{X}\widetilde{\boldsymbol{w}}) + \sum_{i=1}^{p}\sum_{k=1}^{+\infty}\gamma_{ik}\frac{\widetilde{w_{ik}}^2}{2\sigma_k^2} + b_\epsilon\right); \\
P(\sigma_\epsilon^2|\cdot) &\sim IG(\widetilde{a_\epsilon}, \widetilde{b_\epsilon}), \qquad (16) \\
\widetilde{a_\epsilon} &= \frac{n}{2} + \frac{1}{2}\sum_{i=1}^{p}\sum_{k=1}^{+\infty}\gamma_{ik} + a_\epsilon, \\
\widetilde{b_\epsilon} &= \frac{1}{2}(\boldsymbol{E_g} - \boldsymbol{X}\widetilde{\boldsymbol{w}})'\boldsymbol{H}^{-1}(\boldsymbol{E_g} - \boldsymbol{X}\widetilde{\boldsymbol{w}}) + \sum_{i=1}^{p}\sum_{k=1}^{+\infty}\gamma_{ik}\frac{\widetilde{w_{ik}}^2}{2\sigma_k^2} + b_\epsilon.
\end{aligned}$$

- $\sigma_0^2$
  The log conditional density function for $\sigma_0^2$ is given by,

$$\begin{aligned}
log(P(\sigma_0^2|\cdot)) &= C - \frac{1}{2}log|\boldsymbol{H}| - \frac{1}{2\sigma_\epsilon^2}(\boldsymbol{E_g} - \boldsymbol{X}\widetilde{\boldsymbol{w}})'\boldsymbol{H}^{-1}(\boldsymbol{E_g} - \boldsymbol{X}\widetilde{\boldsymbol{w}}) - (a_0 + 1)log(\sigma_0^2) - \frac{b_0}{\sigma_0^2}, \\
\boldsymbol{H} &= \boldsymbol{I} + \sigma_0^2\boldsymbol{K}. \qquad (17)
\end{aligned}$$

  Because the conditional density function (17) is of an unknown distribution, Metropolis-Hastings algorithm is needed to generate posterior samples for $\sigma_0^2$. For improved mixing property, we will re-parametrize $\sigma_0^2$ to $h^2 = \frac{\sigma_0^2}{\sigma_0^2+1}$, such that $h^2$ has domain $[0, 1]$. Based on the probability density function for change of variables $P(h^2|\cdot) = \frac{d\sigma_0^2(h^2)}{dh^2}P(\sigma_0^2|\cdot)$, the log conditional density function for $h^2$ is given by

$$log(P(h^2|\cdot)) = log(\sigma_0^2(h^2)|\cdot) - 2log(1 - h^2), \qquad (18)$$

  where $log(\sigma_0^2(h^2)|\cdot)$ is given by (17) with $\sigma_0^2(h^2) = \frac{h^2}{1-h^2}$. In the MCMC sampling, we will first sample $h^2$ based on (18) and then obtain the $\sigma_0^2$ sample from $\sigma_0^2 = \frac{h^2}{1-h^2}$.

- The above conditional posterior density functions will be used in the Gibbs sampling algorithm to generate posterior samples with respect to each parameter in turn. As a result, the average of the posterior samples will be taken as the posterior Bayesian estimate for the corresponding parameter.

##### 1.2.2 Estimate $\boldsymbol{\zeta}$

Recall that the random effect term $\boldsymbol{u}$ can be represented by the random effect-size vector $\boldsymbol{\zeta}$ as in (5). The posterior conditional distribution for $\boldsymbol{\zeta}$ is given by

$$\begin{aligned}
P(\boldsymbol{\zeta}|\cdot) &\propto P(\boldsymbol{E_g}|\boldsymbol{X}, \widetilde{\boldsymbol{w}}, \boldsymbol{\zeta}, \sigma_\epsilon^2) P(\boldsymbol{\zeta}|\sigma_\epsilon^2, \sigma_0^2) \\
&\propto MVN\left(\boldsymbol{E_g};\ \boldsymbol{X}\widetilde{\boldsymbol{w}} + \boldsymbol{X}\boldsymbol{\zeta},\ \sigma_\epsilon^2 \boldsymbol{I}\right) \times MVN\left(\boldsymbol{\zeta};\ 0,\ \frac{\sigma_\epsilon^2 \sigma_0^2}{p}\boldsymbol{I}\right) \\
&\propto \exp\left\{-\frac{1}{2}\left[-\frac{2}{\sigma_\epsilon^2}\boldsymbol{\zeta}'\boldsymbol{X}'(\boldsymbol{E_g} - \boldsymbol{X}\widetilde{\boldsymbol{w}}) + \frac{1}{\sigma_\epsilon^2}\boldsymbol{\zeta}'\boldsymbol{X}'\boldsymbol{X}\boldsymbol{\zeta} + \frac{p}{\sigma_\epsilon^2\sigma_0^2}\boldsymbol{\zeta}'\boldsymbol{\zeta}\right]\right\} \\
&\propto \exp\left\{-\frac{1}{2}\left[\boldsymbol{\zeta}'\left(\frac{1}{\sigma_\epsilon^2}\boldsymbol{X}'\boldsymbol{X} + \frac{p}{\sigma_\epsilon^2\sigma_0^2}\right)\boldsymbol{\zeta} - 2\boldsymbol{\zeta}'\frac{\boldsymbol{X}'(\boldsymbol{E_g} - \boldsymbol{X}\widetilde{\boldsymbol{w}})}{\sigma_\epsilon^2}\right]\right\}; \\
P(\boldsymbol{\zeta}|\cdot) &\sim MVN(\mu_\zeta, \Sigma_\zeta), \\
\mu_\zeta &= \frac{\boldsymbol{X}'(\boldsymbol{E_g} - \boldsymbol{X}\widetilde{\boldsymbol{w}})}{\sigma_\epsilon^2}\Sigma_\zeta = \frac{\sigma_0^2}{p}\boldsymbol{X}'(\boldsymbol{E_g} - \boldsymbol{X}\widetilde{\boldsymbol{w}})\boldsymbol{H}^{-1}, \qquad (19) \\
\Sigma_\zeta &= \left(\frac{1}{\sigma_\epsilon^2}\boldsymbol{X}'\boldsymbol{X} + \frac{p}{\sigma_\epsilon^2\sigma_0^2}\right)^{-1} = \frac{\sigma_\epsilon^2\sigma_0^2}{p}\left(\sigma_0^2\boldsymbol{K} + \boldsymbol{I}\right)^{-1} = \frac{\sigma_\epsilon^2\sigma_0^2}{p}\boldsymbol{H}^{-1};\ \boldsymbol{K} = \frac{\boldsymbol{X}'\boldsymbol{X}}{p},\ \boldsymbol{H} = \sigma_0^2\boldsymbol{K} + \boldsymbol{I}.
\end{aligned}$$

Instead of sampling $\boldsymbol{\zeta}$ in the MCMC algorithm, we take the Row-Blackwell approximation for the conditional posterior mean of $\boldsymbol{\zeta}$ as the posterior estimate. That is,

$$\widehat{\boldsymbol{\zeta}} = \frac{1}{p}\boldsymbol{X}'\frac{1}{M}\sum_{m=1}^{M}(\sigma_0^2)^{(m)}(\boldsymbol{E_g} - \boldsymbol{X}\widetilde{\boldsymbol{w}}^{(\boldsymbol{m})})(\boldsymbol{H}^{(m)})^{-1}, \qquad (20)$$

where $M$ denotes the total number of MCMC iterations and $(m)$ denotes the corresponding sample value for the $m$th MCMC iteration.

##### 1.2.3 Estimate cis-eQTL Effect-sizes $\boldsymbol{w}$

Therefore, from the MCMC sampling results and the posterior estimate $\widehat{\boldsymbol{\zeta}}$ given by (20), the Bayesian posterior estimate for the cis-eQTL effect-sizes $\widehat{\boldsymbol{w}}$ will be given by

$$\widehat{\boldsymbol{w}} = \widehat{\widetilde{\boldsymbol{w}}} + \widehat{\boldsymbol{\zeta}}. \qquad (21)$$

To improve the computational efficiency of MCMC sampling, the previous DPR paper [1] utilized multiple techniques, e.g., approximating the infinite normal mixture by a truncated normal mixture with $K$ normal components as in [3], implementing an eigen decomposition for $\boldsymbol{K} = \boldsymbol{UDU}'$ and transfer the gene expression levels and genotype as $\boldsymbol{U}'\boldsymbol{E_g}, \boldsymbol{U}'\boldsymbol{X}$ such that the random effects become independent across samples, and taking the random scan Gibbs sampling algorithm [7, 8] to prioritize a set of $500$ cis-eQTL that have the most significant marginal p-values for associated with $\boldsymbol{E_g}$. However, the MCMC sampling still takes ~10 hours to run $50,000$ iterations per gene ($p \approx 10,000$) for the ROS/MAP data with $n = 499$ samples. Thus, we take the variational inference algorithm [1, 3, 9, 10] to fit the nonparametric Bayesian model (4), and then obtain the Bayesian estimates for cis-eQTL effect-sizes $\boldsymbol{w}$.

##### 1.3 Variational Inference Algorithm

Besides the computational expensive MCMC algorithm, the variational inference algorithm [1, 3, 9, 10] provides an alternative, deterministic methodology for approximating likelihoods and posterior density functions for a Bayesian model such as (4). Following the derivations by [1, 3], we will use a particular class of variational methods known as mean-field methods, which is based on optimizing Kullback-Leibler (KL) divergence with respect to a variational distribution.

Let $\boldsymbol{\theta} = \{\widetilde{\boldsymbol{w}}, \boldsymbol{\gamma}, \boldsymbol{u}, \boldsymbol{\nu}, \xi, \sigma_\epsilon^2, \sigma_0^2, \sigma_1^2, \cdots, \sigma_k^2, \cdots\}$ denote the parameters of interest from model (4). Let $q(\boldsymbol{\theta}) = \prod_j q(\theta_j)$ denote a variational distribution that assumes independence among parameters $\{\theta_j\}$ and approximates the joint posterior distribution for $\boldsymbol{\theta}$. We aim to identify a variational distribution such that the KL divergence between the variational distribution $q(\boldsymbol{\theta})$ and the joint posterior distribution $P(\boldsymbol{\theta}|\boldsymbol{E_g}, \boldsymbol{X}, \boldsymbol{K})$ is minimized. In particular,

$$\begin{aligned} KL(q(\boldsymbol{\theta})|P(\boldsymbol{\theta}|\boldsymbol{E_g}, \boldsymbol{X}, \boldsymbol{K})) &= E_{q(\boldsymbol{\theta})}\left[log\left(\frac{q(\boldsymbol{\theta})}{P(\boldsymbol{\theta}|\boldsymbol{E_g}, \boldsymbol{X}, \boldsymbol{K})}\right)\right] \qquad (22) \\ &= E_{q(\boldsymbol{\theta})}\left[log(q(\boldsymbol{\theta}))\right] - E_{q(\boldsymbol{\theta})}\left[log(P(\boldsymbol{\theta}, \boldsymbol{E_g}, \boldsymbol{X}, \boldsymbol{K}))\right] + log(P(\boldsymbol{E_g}, \boldsymbol{X}, \boldsymbol{K})). \end{aligned}$$

Since $P(\boldsymbol{E_g}, \boldsymbol{X}, \boldsymbol{K})$ in (22) is independent of $q(\boldsymbol{\theta})$, minimizing the KL divergence (22) is equivalent to maximizing the evidence lower bound (ELBO),

$$ELBO = E_{q(\boldsymbol{\theta})}\left[log(P(\boldsymbol{\theta}, \boldsymbol{E_g}, \boldsymbol{X}, \boldsymbol{K}))\right] - E_{q(\boldsymbol{\theta})}\left[log(q(\boldsymbol{\theta}))\right]. \qquad (23)$$

Because of the independence among $\{q(\theta_j)\}$, a gradient ascent algorithm [11] can be taken to maximize $(23)$ by maximizing the ELBO with respect to each $q(\theta_j)$ in turn. We can derive the partial derivative of ELBO with respect to $q(\theta_j)$ as follows,

$$\begin{aligned} \frac{\partial ELBO}{\partial q(\theta_j)} &= \frac{\partial\left(\int q(\theta_j)E_{q(-\theta_j)}[log(P(\boldsymbol{\theta}, \boldsymbol{E_g}, \boldsymbol{X}, \boldsymbol{K}))]d\theta_j - \int q(\theta_j)log(q(\theta_j))d\theta_j - \int q(\theta_j)C_{-j}d\theta_j\right)}{\partial q(\theta_j)} \\ &= E_{q(-\theta_j)}[log(P(\boldsymbol{\theta}, \boldsymbol{E_g}, \boldsymbol{X}, \boldsymbol{K}))] - log(q(\theta_j)) - C_{-j}, \qquad (24) \end{aligned}$$

where $E_{q(\boldsymbol{\theta})}\left[log(q(\boldsymbol{\theta}))\right] = E_{q(\boldsymbol{\theta})}\left[\sum_j log(q(\theta_j))\right] = E_{q(\boldsymbol{\theta})}\left[log(q(\theta_j))\right] + E_{q(\boldsymbol{\theta})}\left[\sum_{l \neq j} log(q(\theta_l))\right] = E_{q(\boldsymbol{\theta})}\left[log(q(\theta_j))\right] + \int q(\theta_j)C_{-j}d\theta_j$, and $C_{-j}$ is a constant free of $q(\theta_j)$. By setting the partial derivative (24) equal to zero, we obtain the following variational distribution for $\theta_j$,

$$q(\theta_j) \propto \exp\left\{E_{q(\theta_{-j})}[log(P(\boldsymbol{\theta}, \boldsymbol{E_g}, \boldsymbol{X}, \boldsymbol{K}))]\right\} \propto \exp\left\{E_{q(\theta_{-j})}[log(P(\theta_j|\theta_{-j}, \boldsymbol{E_g}, \boldsymbol{X}, \boldsymbol{K}))]\right\}, \qquad (25)$$

where $\theta_{-j}$ denotes other parameters excluding $\theta_j$, and $q(\theta_{-j})$ denotes the variational distribution for $\theta_{-j}$.

As a result, the optimal variational distributions per $\theta_j$ can be obtained by the gradient ascent algorithm with a partial derivative quantity (24). The ELBO quantity (23) will be used to check convergence of the gradient ascent algorithm. Then the posterior mean of the corresponding optimal variational distribution $q(\theta_j)$ from the last iteration will be taken as the Bayesian estimator for $\theta_j$.

##### 1.3.1 Joint Posterior Distribution for $\boldsymbol{\theta}$

Because the ELBO quantity (23) is difficult to compute if the variational distributions are of unknown distributions, we will derive the variational inference algorithm without integrating out the random effect term $\boldsymbol{u}$. For computational convenience, we transform the model (4) as follows,

$$\mathbf{U'E_g} \quad = \quad \mathbf{U'X\widetilde{w} + U'u} + \varepsilon; \ \boldsymbol{\eta} = \mathbf{U'u} \sim \mathbf{N(0, \sigma_\epsilon^2\sigma_0^2 D)}; \ \mathbf{K} = \frac{\boldsymbol{XX'}}{\boldsymbol{p}} = \mathbf{U'DU}, \qquad (26)$$

where $\boldsymbol{D}$ is a diagonal matrix and prior distributions for $\{\widetilde{\boldsymbol{w}}, \sigma_0^2, \sigma_\epsilon^2\}$ are the same as in model (4).

With indicator vectors $\boldsymbol{\gamma}$, the joint conditional posterior function for $\boldsymbol{\theta}$ is given by,

$$\begin{aligned} P(\boldsymbol{\theta}|\boldsymbol{E_g}, \boldsymbol{X}, \boldsymbol{K}) = P(\widetilde{\boldsymbol{w}}, \boldsymbol{\gamma}, \boldsymbol{\eta}, \boldsymbol{\nu}, \xi, \sigma_\epsilon^2, \sigma_0^2, \sigma_1^2, \cdots, \sigma_k^2, \cdots|\boldsymbol{E_g}, \boldsymbol{X}, \boldsymbol{K}) \propto \\ P(\boldsymbol{U'E_g}|\boldsymbol{U'X}, \boldsymbol{D}, \widetilde{\boldsymbol{w}}, \boldsymbol{\eta}, \sigma_\epsilon^2) P(\boldsymbol{w}|\boldsymbol{\gamma}, \boldsymbol{\nu}, \sigma_1^2, \cdots, \sigma_k^2, \cdots) P(\boldsymbol{\eta}|\sigma_\epsilon^2, \sigma_0^2, \boldsymbol{D}) \\ \left(\prod_{k=0}^{+\infty} P(\sigma_k^2|a_k, b_k)\right) P(\sigma_\epsilon^2|a_\epsilon, b_\epsilon) P(\boldsymbol{\gamma}|\boldsymbol{\nu}) P(\boldsymbol{\nu}|\xi) P(\xi|a_\xi, b_\xi). \end{aligned} \qquad (27)$$

Consequently, the log joint posterior density function is given by

$$\begin{aligned} log(P(\widetilde{\boldsymbol{w}}, \boldsymbol{\gamma}, \boldsymbol{\eta}, \boldsymbol{\nu}, \xi, \sigma_\epsilon^2, \sigma_0^2, \sigma_1^2, \cdots, \sigma_k^2, \cdots|\boldsymbol{E_g}, \boldsymbol{X}, \boldsymbol{K})) = \\ C - \tfrac{n}{2}log(\sigma_\epsilon^2) - \tfrac{1}{2\sigma_\epsilon^2}(\boldsymbol{U'E_g} - \boldsymbol{U'X}\widetilde{\boldsymbol{w}} - \boldsymbol{\eta})'(\boldsymbol{U'E_g} - \boldsymbol{U'X}\widetilde{\boldsymbol{w}} - \boldsymbol{\eta}) + \\ \textstyle\sum_{i=1}^{p}\sum_{k=1}^{+\infty}\gamma_{ik}\left[-\tfrac{1}{2}log(\sigma_\epsilon^2) - \tfrac{1}{2}log(\sigma_k^2) - \tfrac{\widetilde{w_{ik}}^2}{2\sigma_\epsilon^2\sigma_k^2}\right] - \tfrac{n}{2}log(\sigma_\epsilon^2) - \tfrac{n}{2}log(\sigma_0^2) - \tfrac{1}{2}\boldsymbol{\eta}'(\sigma_\epsilon^2\sigma_0^2\boldsymbol{D})^{-1}\boldsymbol{\eta} + \\ \textstyle\sum_{k=0}^{+\infty}\left[-(a_k+1)log(\sigma_k^2) - \tfrac{b_k}{\sigma_k^2}\right] - (a_\epsilon+1)log(\sigma_\epsilon^2) - \tfrac{b_\epsilon}{\sigma_\epsilon^2} + \\ \textstyle\sum_{i=1}^{p}\sum_{k=1}^{+\infty}\left[\gamma_{ik}\left(log(\nu_k) + \sum_{l=0}^{k-1}log(1-\nu_l)\right) + (1-\gamma_{ik})log\left(1 - \nu_k\prod_{l=0}^{k-1}(1-\nu_l)\right)\right] + \\ \textstyle\sum_{k=0}^{+\infty}[(\xi-1)log(1-\nu_k) + log(\xi)] + (a_\xi - 1)log(\xi) - b_\xi\xi, \end{aligned} \qquad (28)$$

where $C$ denotes a normalization constant that is free of parameters.

##### 1.3.2 Variational Distribution for $\theta_j$

Based on the above log joint posterior density function (28), the following variational distributions are derived with respect to each $\theta_j$.

- $\widetilde{w_{ik}}$ **and** $\gamma_{ik}$

$$\begin{aligned}
log(q(\widetilde{w_{ik}}, \gamma_{ik})) &= C - \frac{1}{2}E[\sigma_\epsilon^{-2}]\boldsymbol{x_i'UU'x_i}(\widetilde{w}_{ik}^2 + 2\widetilde{w}_{ik}E[\widetilde{w}_{i(-k)}]) + \\
& E[\sigma_\epsilon^{-2}]\boldsymbol{x_i'U}\left(\boldsymbol{U'E_g} - \sum_{l\neq i}\left(\boldsymbol{U'x_l}E[\widetilde{w_l}]\right) - E[\boldsymbol{\eta}]\right)\widetilde{w_{ik}} + \\
& \gamma_{ik}\left[-\frac{1}{2}E[log(\sigma_\epsilon^2)] - \frac{1}{2}E[log(\sigma_k^2)] - \frac{1}{2}E[\sigma_\epsilon^{-2}]E[\sigma_k^{-2}]\widetilde{w_{ik}}^2\right] + \\
& \gamma_{ik}\left(E[log(\nu_k)] + \sum_{l=0}^{k-1}E[log(1-\nu_l)]\right) + \\
& (1-\gamma_{ik})E\left[log\left(1 - \nu_k\prod_{l=0}^{k-1}(1-\nu_l)\right)\right], \qquad (29)
\end{aligned}$$

where $\widetilde{w}_{i(-k)} = \sum_{j\neq k}\widetilde{w}_{ij}$.

Then $q(\widetilde{w_{ik}}|\gamma_{ik} = 1)$ is given by

$$\begin{aligned}
q(\widetilde{w_{ik}}|\gamma_{ik} = 1) &= C - \frac{1}{2}E[\sigma_\epsilon^{-2}]\boldsymbol{x_i'UU'x_i}\widetilde{w_{ik}}^2 - \frac{1}{2}E[\sigma_\epsilon^{-2}]E[\sigma_k^{-2}]\widetilde{w_{ik}}^2 + \\
& E[\sigma_\epsilon^{-2}]\boldsymbol{x_i'U}\left(\boldsymbol{U'E_g} - \sum_{l\neq i}(\boldsymbol{U'x_l}E[\widetilde{w_l}]) - E[\boldsymbol{\eta}] - \boldsymbol{U'x_i}E[\widetilde{w}_{i(-k)}]\right)\widetilde{w_{ik}} \\
&= C - \frac{E[\sigma_\epsilon^{-2}]}{2}\left(\boldsymbol{x_i'U'Ux_i} - E[\sigma_k^{-2}]\right)\widetilde{w_{ik}}^2 + \\
& E[\sigma_\epsilon^{-2}]\boldsymbol{x_i'U'}\left(\boldsymbol{UE_g} - \sum_{l\neq i}(\boldsymbol{U'x_l}E[\widetilde{w_l}]) - E[\boldsymbol{\eta}] - \boldsymbol{U'x_i}E[\widetilde{w}_{i(-k)}]\right)\widetilde{w_{ik}} \\
q(\widetilde{w_{ik}}|\gamma_{ik} = 1) &\sim N(m_{ik}, s_{ik}^2), \\
m_{ik} &= \frac{\boldsymbol{x_i'U}(\boldsymbol{U'E_g} - \sum_{l\neq i}\boldsymbol{U'x_l}E[\widetilde{w_l}] - E[\boldsymbol{\eta}] - \boldsymbol{U'x_i}E[\widetilde{w}_{i(-k)}])}{\boldsymbol{x_i'U'Ux_i} + E[\sigma_k^{-2}]}, \\
s_{ik}^2 &= \frac{1}{E[\sigma_\epsilon^{-2}](\boldsymbol{x_i'UU'x_i} + E[\sigma_k^{-2}])}. \qquad (30)
\end{aligned}$$

Similar to the derivation for (12), after integrating out $\widetilde{w_{ik}}$ from (29), the variational probability for $(\gamma_{ik} = 1)$ is given by

$$\begin{aligned}
& q(\gamma_{ik} = 1) = \phi_{ik} = \int q(\widetilde{w_{ik}}, \gamma_{ik} = 1)d\widetilde{w_{ik}} \propto \\
& \exp\left\{\frac{m_{ik}^2}{2s_k^2} + log(s_k) - E[log(\sigma_\epsilon)] - E[log(\sigma_k)] + E[log(\nu_k)] + \sum_{l=0}^{k-1}E[log(1-\nu_l)]\right\}. \qquad (31)
\end{aligned}$$

- $\nu_k$

$$log(q(\nu_k)) = C + \sum_{i=1}^{p} E[\gamma_{ik}]log(\nu_k) + \sum_{i=1}^{p}\sum_{l=k+1}^{+\infty} E[\gamma_{ik}]log(1-\nu_k) + (E[\xi]-1)log(1-\nu_k),$$
$$q(\nu_k) \sim Beta(\kappa_k, \xi_k),\ \kappa_k = \sum_{i=1}^{p} E[\gamma_{ik}] + 1,\ \xi_k = \sum_{i=1}^{p}\sum_{l=k+1}^{+\infty} E[\gamma_{il}] + E[\xi]. \quad (32)$$

- $\sigma_k^2, k > 0$

$$log(q(\sigma_k^2)) = C - \left(\frac{1}{2}\sum_{i=1}^{p} E[\gamma_{ik}] + a_k + 1\right) log(\sigma_k^2) - \left(\frac{\sum_{i=1}^{p} E[\gamma_{ik}\widetilde{w_{ik}}^2]E[\sigma_\epsilon^{-2}]}{2} + b_k\right)\frac{1}{\sigma_k^2};$$
$$q(\sigma_k^2) \sim IG(\widetilde{a_k}, \widetilde{b_k}),\ \widetilde{a_k} = \frac{1}{2}\sum_{i=1}^{p} E[\gamma_{ik}] + a_k,\ \widetilde{b_k} = \frac{1}{2}\sum_{i=1}^{p} E[\gamma_{ik}\widetilde{w_{ik}}^2]E[\sigma_\epsilon^{-2}] + b_k. \quad (33)$$

- $\xi$

$$log(q(\xi)) = C + \left(a_\xi + \sum_{k=0}^{+\infty} 1_k - 1\right) log(\xi) - \left(b_\xi - \sum_{k=0}^{+\infty} E[log(1-\nu_k)]\right)\xi;$$
$$q(\xi) \sim Gamma(\widetilde{a_\xi}, \widetilde{b_\xi}),\ \widetilde{a_\xi} = a_\xi + \sum_{k=0}^{+\infty} 1_k,\ \widetilde{b_\xi} = b_\xi - \sum_{k=0}^{+\infty} E[log(1-\nu_k)]. \quad (34)$$

- $\sigma_\epsilon^2$

$$\begin{aligned} log(q(\sigma_\epsilon^2)) &= C - log(\sigma_\epsilon^2)\left(\frac{n}{2} + \frac{1}{2}\sum_{i=1}^{p}\sum_{k=1}^{+\infty} E[\gamma_{ik}] + a_\epsilon + 1\right) - \\ &\quad \frac{1}{2\sigma_\epsilon^2}(\boldsymbol{U'E_g} - \boldsymbol{U'XE}[\widetilde{\boldsymbol{w}}] - E[\boldsymbol{\eta}])'(\boldsymbol{U'E_g} - \boldsymbol{U'XE}[\widetilde{\boldsymbol{w}}] - E[\boldsymbol{\eta}]) - \quad (35) \\ &\quad \frac{1}{2\sigma_\epsilon^2}\sum_{i=1}^{p}\sum_{k=1}^{+\infty} E[\gamma_{ik}\widetilde{w_{ik}}^2]E[\sigma_k^{-2}] - \frac{1}{2\sigma_\epsilon^2}\sum_{i=1}^{p}\frac{E[\eta_i^2]E[\sigma_0^{-2}]}{D_{ii}} - \frac{1}{\sigma_\epsilon^2}b_\epsilon; \\ q(\sigma_\epsilon^2) &\sim IG(\widetilde{a_\epsilon}, \widetilde{b_\epsilon}), \quad (36) \\ \widetilde{a_\epsilon} &= \frac{n}{2} + \frac{1}{2}\sum_{i=1}^{p}\sum_{k=1}^{+\infty} E[\gamma_{ik}] + a_\epsilon, \\ \widetilde{b_\epsilon} &= \frac{1}{2}(\boldsymbol{U'E_g} - \boldsymbol{U'XE}[\widetilde{\boldsymbol{w}}] - E[\boldsymbol{\eta}])'(\boldsymbol{U'E_g} - \boldsymbol{U'XE}[\widetilde{\boldsymbol{w}}] - E[\boldsymbol{\eta}]) + \\ &\quad \frac{1}{2}\left(\sum_{i=1}^{p}\sum_{k=1}^{+\infty} E[\gamma_{ik}\widetilde{w_{ik}}^2]E[\sigma_k^{-2}] + \sum_{i=1}^{p}\frac{E[\eta_i^2]E[\sigma_0^{-2}]}{D_{ii}}\right) + b_\epsilon. \end{aligned}$$

- $\sigma_0^2$

$$\begin{aligned} log(q(\sigma_0^2)) &= C - \frac{n}{2}log(\sigma_0^2) - \frac{1}{2\sigma_0^2}\sum_{i=1}^{p}\frac{E[\eta_i]^2 E[\sigma_\epsilon^{-2}]}{D_{ii}} - (a_0+1)log(\sigma_0^2) - \frac{b_0}{\sigma_0^2}, \\ q(\sigma_0^2) &\sim IG(\widetilde{a_0},\ \widetilde{b_0}),\ \widetilde{a_0} = \frac{n}{2} + a_0,\ \widetilde{b_0} = \frac{1}{2}\sum_{i=1}^{p}\frac{E[\eta_i]^2 E[\sigma_\epsilon^{-2}]}{D_{ii}} + b_0 \end{aligned} \tag{37}$$

- $\boldsymbol{\eta}$

$$\begin{aligned} log(q(\boldsymbol{\eta})) &= C - \frac{E[\sigma_\epsilon^{-2}]}{2}(\boldsymbol{U'E_g} - \boldsymbol{U'XE}[\widetilde{\boldsymbol{w}}] - \boldsymbol{\eta})'(\boldsymbol{U'E_g} - \boldsymbol{U'XE}[\widetilde{\boldsymbol{w}}] - \boldsymbol{\eta}) - \\ &\quad \frac{1}{2}\boldsymbol{\eta}'E[(\sigma_\epsilon^2\sigma_0^2\boldsymbol{D})^{-1}]\boldsymbol{\eta}; \\ q(\boldsymbol{\eta}) &\sim MVN(\boldsymbol{\mu_\eta},\ \boldsymbol{\Sigma_\eta}), \\ \boldsymbol{\mu_\eta} &= (E[\sigma_0^{-2}]\boldsymbol{D}^{-1} + \boldsymbol{I})^{-1}(\boldsymbol{U'E_g} - \boldsymbol{U'XE}[\widetilde{\boldsymbol{w}}]), \\ \boldsymbol{\Sigma_\eta} &= \frac{(E[\sigma_0^{-2}]\boldsymbol{D}^{-1} + \boldsymbol{I})^{-1}}{E[\sigma_\epsilon^{-2}]}; \end{aligned} \tag{38}$$

where $\boldsymbol{\Sigma_\eta}$ is a diagonal matrix. That is, $\{q(\eta_i); i = 1, \cdots, p\}$ are independent of each other.

- Evaluations of the expectations in the above variational distributions:

$$\begin{aligned} E[\boldsymbol{\eta}] &= \boldsymbol{\mu_\eta}; \\ E[\gamma_{ik}] &= \phi_{ik},\ i = 1, \cdots, p,\ k = 1, 2, \cdots; \\ E[\gamma_i \widetilde{w_i}^2] &= \sum_k \phi_{ik}(m_{ik}^2 + s_{ik}^2);\ E[\widetilde{w_i}] = \sum_k \phi_{ik} m_{ik};\ i = 1, \cdots, p \\ E[log(\nu_k)] &= \psi(\kappa_k) - \psi(\kappa_k + \xi_k);\ E[log(1-\nu_k)] = \psi(\xi_k) - \psi(\kappa_k + \xi_k);\ k = 0, 1, \cdots \\ E[log(\sigma_k)] &= \frac{1}{2}(log(\widetilde{b_k}) - \psi(\widetilde{a_k}));\ E[\sigma_k^{-2}] = \frac{\widetilde{a_k}}{\widetilde{b_k}};\ k = 0, 1, \cdots \\ E[log(\sigma_\epsilon)] &= \frac{1}{2}(log(\widetilde{b_\epsilon}) - \psi(\widetilde{a_\epsilon}));\ E[\sigma_\epsilon^{-2}] = \frac{\widetilde{a_\epsilon}}{\widetilde{b_\epsilon}}; \\ E[\log(\xi)] &= \psi(\widetilde{a_\xi}) - log(\widetilde{b_\xi});\ E[\xi] = \frac{\widetilde{a_\xi}}{\widetilde{b_\xi}}; \end{aligned}$$

where $\psi(x) = \frac{\Gamma'(x)}{\Gamma(x)}$ is the digamma function and the expectations are taken with respect to each variational distribution $q(\theta_j)$.

##### 1.3.3 Evaluate the ELBO

Recall the formula (23) for ELBO, we will need to calculate the expectation quantities $E_{q(\boldsymbol{\theta})}\left[log(q(\boldsymbol{\theta}))\right]$ and $E_{q(\boldsymbol{\theta})}\left[log(P(\boldsymbol{\theta}, \boldsymbol{E_g}, \boldsymbol{X}, \boldsymbol{K}))\right]$. In particular,

$$\begin{aligned} E_{q(\boldsymbol{\theta})}\left[log(q(\boldsymbol{\theta}))\right] &= \sum_j E_{q(\boldsymbol{\theta})}\left[log(q(\theta_j))\right] \\ &= \sum_{i=1}^{p}\sum_{k=1}^{+\infty} E_{q(\widetilde{w_i},\gamma_i))}[log(q(\widetilde{w_i},\gamma_i))] + \sum_{i=1}^{p} E_{q(\eta_i)}[log(q(\eta_i))] + \\ &\quad \sum_{k=1}^{+\infty} E_{q(\nu_k)}[log(q(\nu_k))] + \sum_{k=1}^{+\infty} E_{q(\sigma_k^2)}[log(q(\sigma_k^2))] + \\ &\quad E_{q(\sigma_\epsilon^2)}[log(q(\sigma_\epsilon^2))] + E_{q(\xi)}[log(q(\xi))], \end{aligned} \tag{39}$$

where

$$\begin{aligned} E_{q(\widetilde{w_i},\gamma_i))}[log(q(\widetilde{w_i},\gamma_i))] &= \sum_{k=1}^{+\infty}\phi_{ik}E[log(q(\widetilde{w_i},\gamma_i=1))] = \sum_{k=1}^{+\infty}\phi_{ik}(E[log(q(\widetilde{w_i}|\gamma_i=1)q(\gamma_i=1))]) \\ &= \sum_{k=1}^{+\infty}\phi_{ik}\left(log(\phi_{ik}) + E[log(q(\widetilde{w_i}|\gamma_i=1))]\right) \\ &= \sum_{k=1}^{+\infty}\phi_{ik}\left(log(\phi_{ik}) - \frac{1}{2}log(2\pi s_{ik}^2) - \frac{1}{2}\right); \\ E_{q(\eta_i)}[log(q(\eta_i))] &= -\frac{1}{2}log(2\pi\Sigma_{\eta_{ii}}) - \frac{1}{2},\ i=1,\cdots,p; \end{aligned}$$

$$\begin{aligned} E_{q(\nu_k)}[log(q(\nu_k))] &= log(\Gamma(\kappa_k+\xi_k)) - log(\Gamma(\kappa_k)) - log(\Gamma(\xi_k)) + \\ &\quad (\kappa-1)E[log(\nu_k)] + (\xi_k-1)E[log(1-\nu_k)] \\ &= log(\Gamma(\kappa_k+\xi_k)) - log(\Gamma(\kappa_k)) - log(\Gamma(\xi_k)) + \\ &\quad (\kappa-1)(\psi(\kappa_k)-\psi(\kappa_k+\xi_k)) - (\xi_k-1)(\psi(\kappa_k+\xi_k)-\psi(\xi_k)); \\ E_{q(\sigma_k^2)}[log(q(\sigma_k^2))] &= \widetilde{a}_k log(\widetilde{b}_k) - log(\Gamma(\widetilde{a}_k)) - 2(\widetilde{a}_k+1)E[log(\sigma_k)] - \widetilde{b}_k E[\sigma_k^{-2}] \\ &= \widetilde{a}_k log(\widetilde{b}_k) - log(\Gamma(\widetilde{a}_k)) - (\widetilde{a}_k+1)(log(\widetilde{b}_k)-\psi(\widetilde{a}_k)) - \widetilde{a}_k,\ k=1,2,\cdots; \\ E_{q(\sigma_\epsilon^2)}[log(q(\sigma_\epsilon^2))] &= \widetilde{a}_\epsilon log(\widetilde{b}_\epsilon) - log(\Gamma(\widetilde{a}_\epsilon)) - (\widetilde{a}_\epsilon+1)(log(\widetilde{b}_\epsilon)-\psi(\widetilde{a}_\epsilon)) - \widetilde{a}_\epsilon; \\ E_{q(\xi)}[log(q(\xi))] &= \widetilde{a}_\xi log(\widetilde{b}_\xi) - log(\Gamma(\widetilde{a}_\xi)) + (\widetilde{a}_\xi-1)E[log(\xi)] - \widetilde{b}_\xi E[\xi] \\ &= log(\widetilde{b}_\xi) - log(\Gamma(\widetilde{a}_\xi)) + (\widetilde{a}_\xi-1)\psi(\widetilde{a}_\xi) - \widetilde{a}_\xi. \end{aligned}$$

And

$$E_{q(\boldsymbol{\theta})}\left[log(P(\boldsymbol{\theta}, \boldsymbol{E_g}, \boldsymbol{X}, \boldsymbol{U}, \boldsymbol{D}))\right] \tag{40}$$

$$\begin{aligned} &= E_{q(\boldsymbol{\theta})}\left[log(P(\boldsymbol{E_g}|\boldsymbol{\theta},\boldsymbol{X},\boldsymbol{U})P(\widetilde{w}|\boldsymbol{\gamma},\sigma_\epsilon^2,\sigma_k^2,k=1,\cdots)P(\boldsymbol{\eta}|\boldsymbol{D},\sigma_\epsilon^2,\sigma_0^2)P(\boldsymbol{\gamma}|\boldsymbol{\nu})P(\boldsymbol{\nu}|\xi)P(\xi))\right] \\ &= E_{q(\boldsymbol{\theta})}[log(P(\boldsymbol{E_g}|\boldsymbol{\theta},\boldsymbol{X})] + E_{q(\boldsymbol{\theta})}[log(P(\widetilde{w}|\boldsymbol{\gamma},\sigma_\epsilon^2,\sigma_k^2,k=1,\cdots))] + E_{q(\boldsymbol{\theta})}[log(P(\boldsymbol{\eta}|\boldsymbol{D},\sigma_\epsilon^2,\sigma_0^2))] + \\ &\quad E_{q(\boldsymbol{\theta})}[log(P(\boldsymbol{\gamma}|\boldsymbol{\nu}))] + E_{q(\boldsymbol{\theta})}[log(P(\boldsymbol{\nu}|\xi))] + E_{q(\boldsymbol{\theta})}[log(P(\xi))]. \end{aligned} \tag{41}$$

##### 1.3.4 Gradient Ascent Algorithm

Starting with initial parameter values $\boldsymbol{\theta}_0$, we will iteratively evaluate the above variational distributions and take the variational means as the corresponding parameter values in next iteration. Iterations will stop when the ELBO quantity converges.

The variational means from the last iteration will be taken as the Bayesian estimates for corresponding parameters, e.g., $\widehat{\widetilde{\boldsymbol{w}}}, \widehat{\boldsymbol{\eta}}$. Recall that $\boldsymbol{\eta} = \boldsymbol{U}'\boldsymbol{X}\boldsymbol{\zeta}$, then the Bayesian estimate for $\boldsymbol{\zeta}$ is given by $\widehat{\boldsymbol{\zeta}} = (\boldsymbol{U}'\boldsymbol{X})^{-1}\widehat{\boldsymbol{\eta}}$. Thus, the Bayesian estimate for cis-eQTL effect-sizes is given by $\widehat{\boldsymbol{w}} = \widehat{\widetilde{\boldsymbol{w}}} + \widehat{\boldsymbol{\zeta}}$.

#### 2 Supplemental Figures

**Figure S** 1

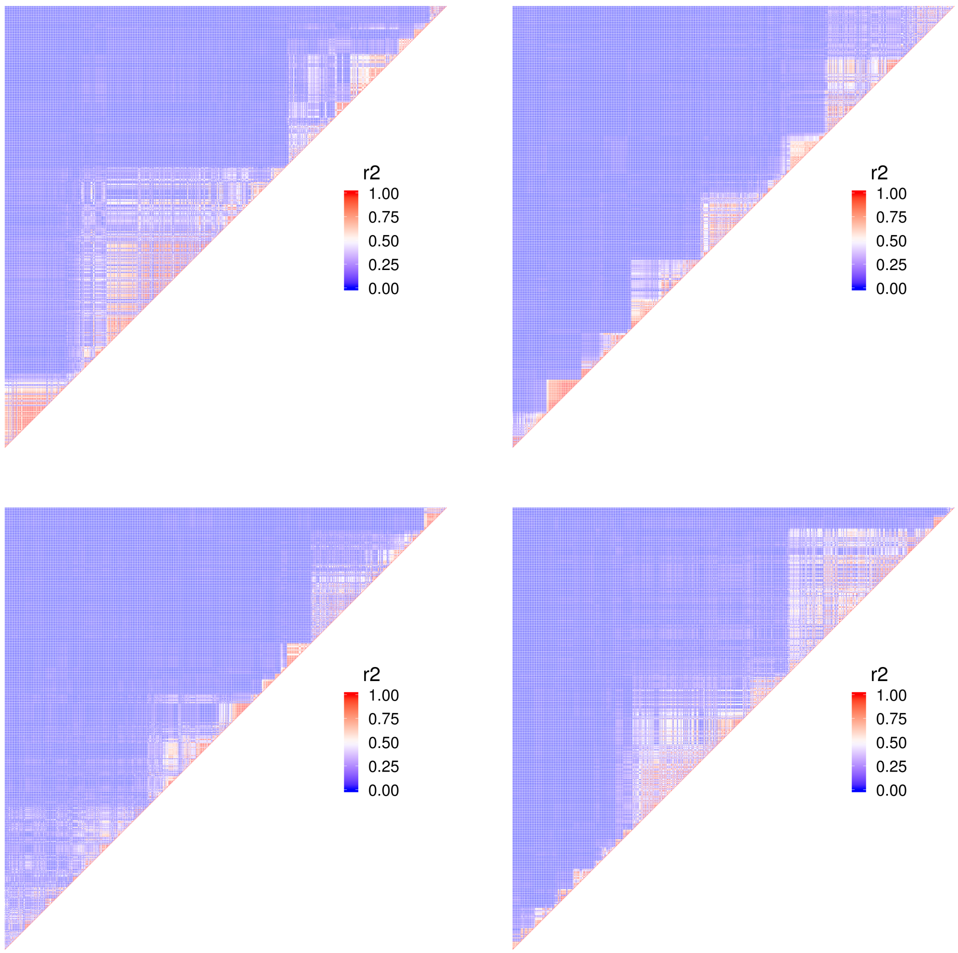


Linkage disequilibrium block structure for *ABCA7*, where each plot represents a non-overlapped region of the genotype data used in our simulations studies.

**Figure S** 2

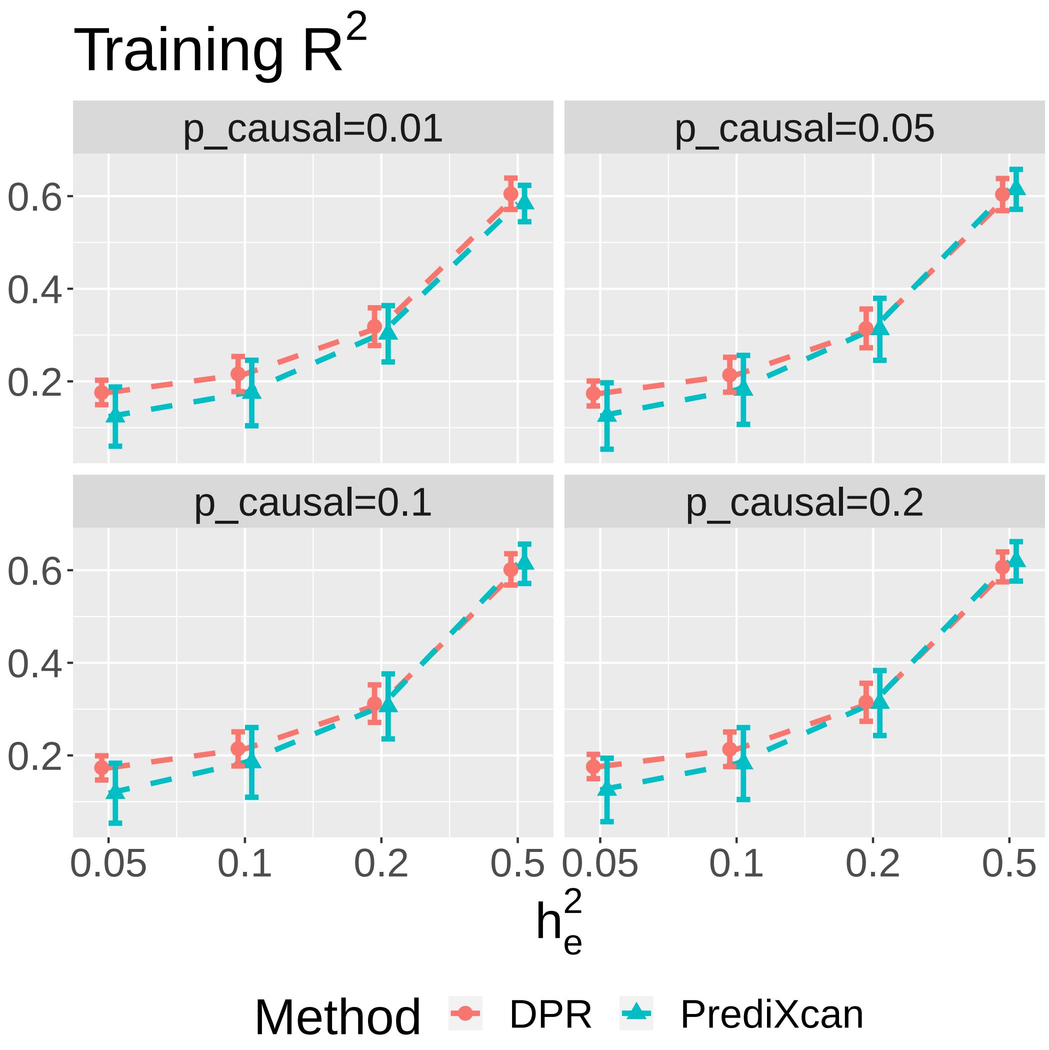


Training $R^2$ by **DPR** and **PrediXcan** under various simulation scenarios with the proportions of true causal SNPs $p_{causal} = (0.01, 0.05, 0.1, 0.2)$ and expression heritability $h_e^2 = (0.05, 0.1, 0.2, 0.5)$.

**Figure S** 3

(A)

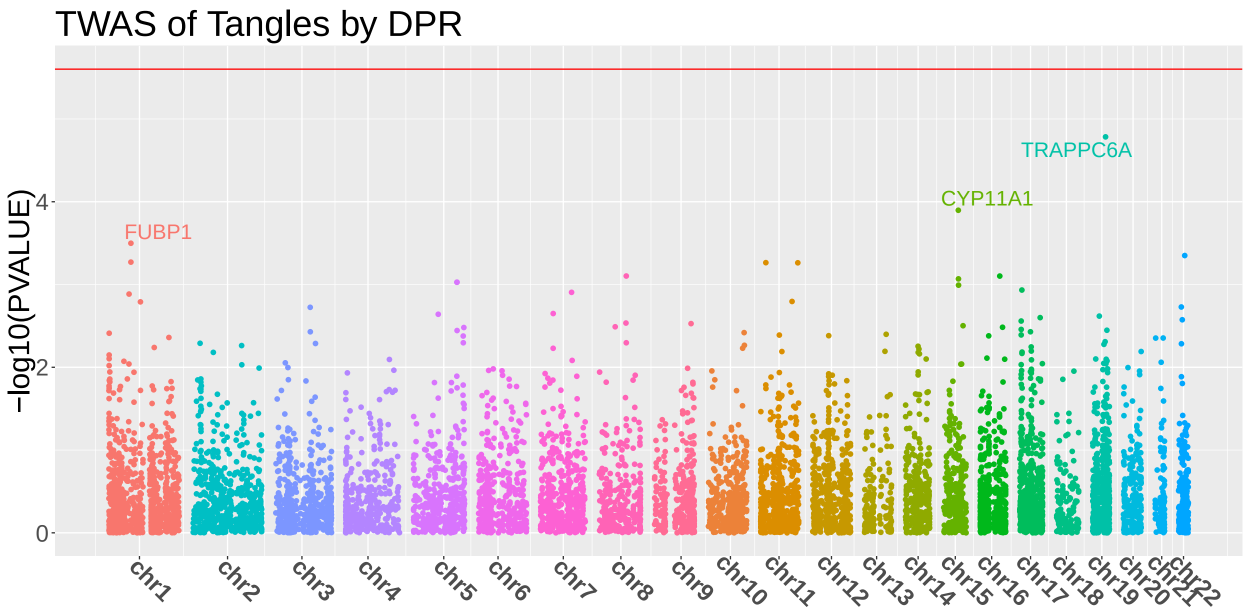


(B)

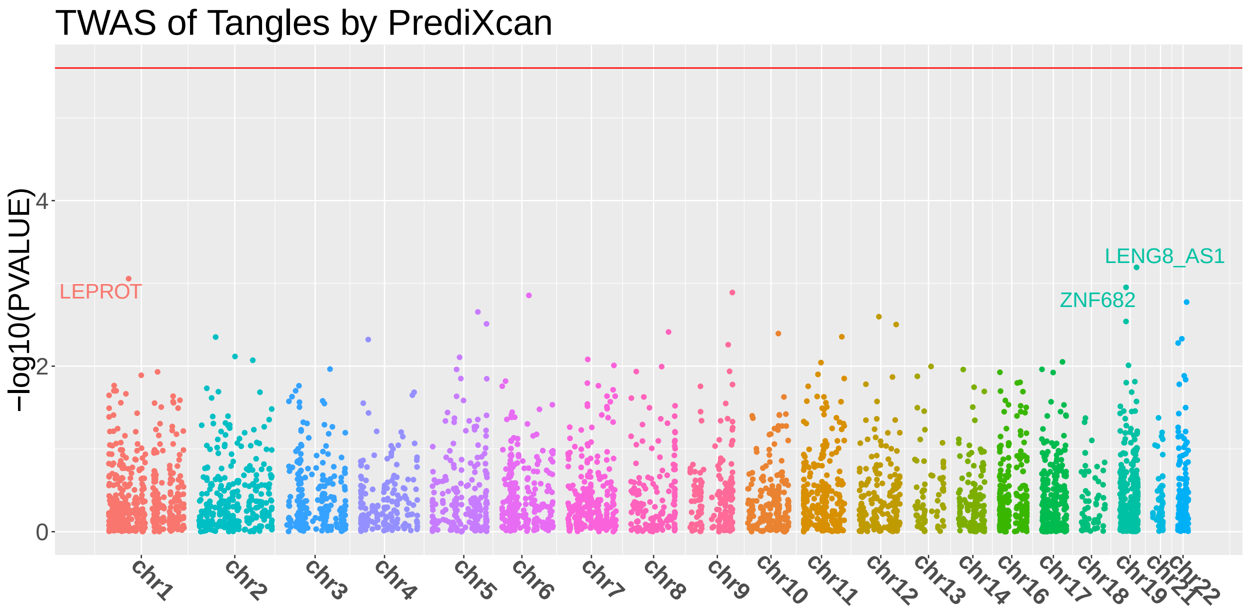


Manhattan plots for TWAS of neurofibrillary tangle density (Tangles) using individual-level ROS/MAP data and imputation models by **DPR (A)** and **PrediXcan (B)**.

**Figure S** 4

(A)

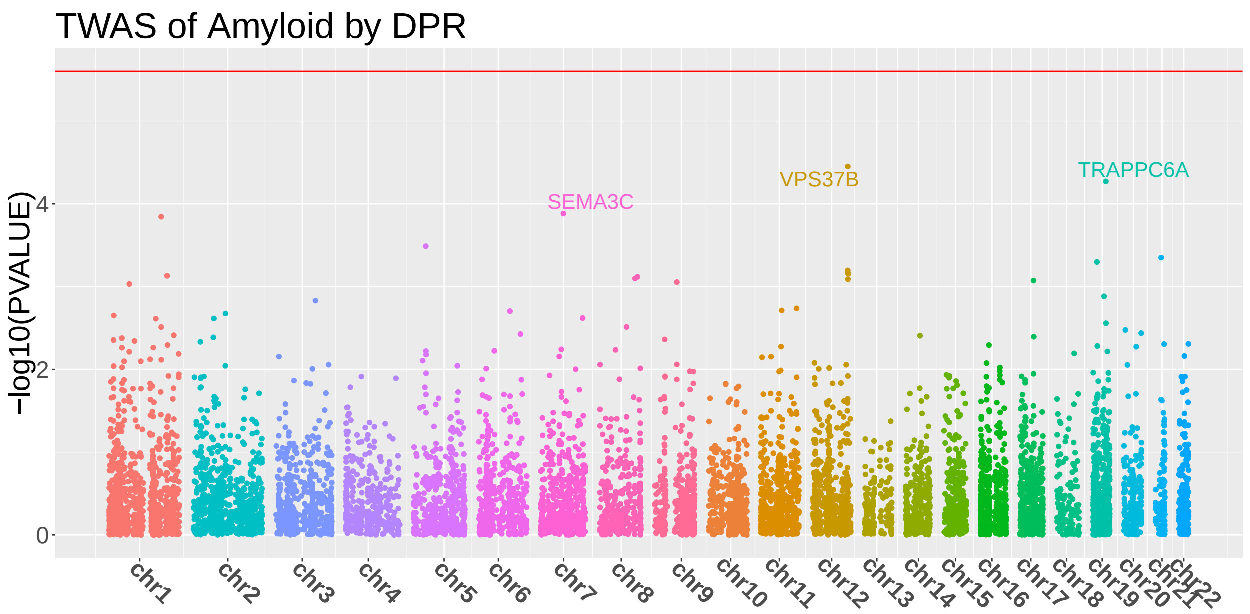


(B)

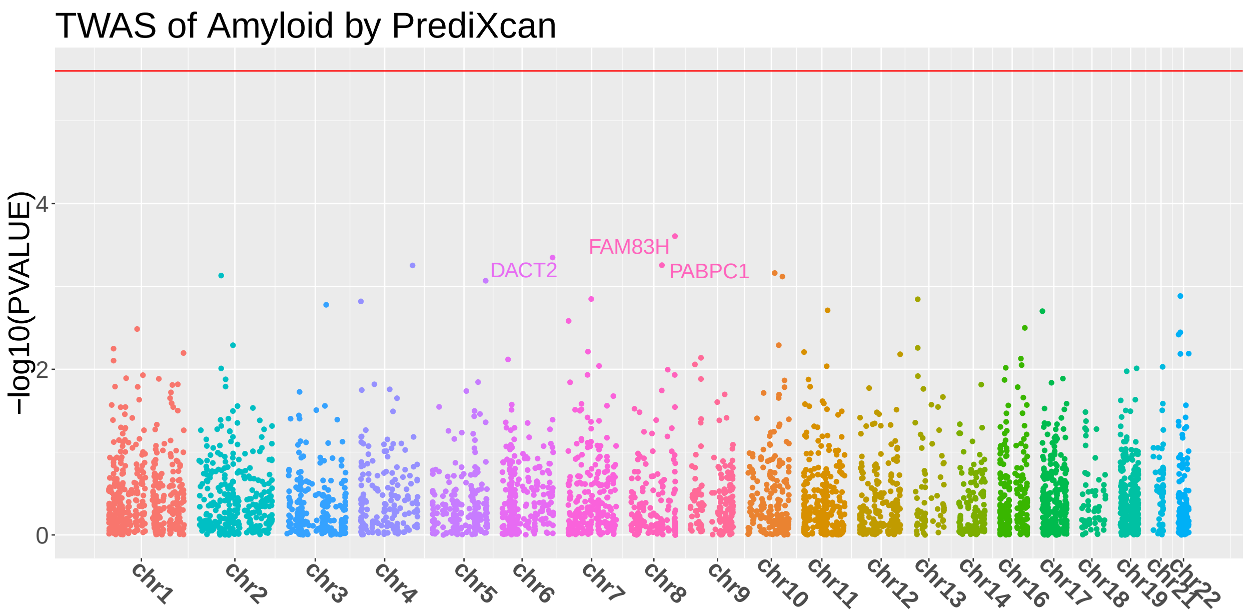


Manhattan plots for TWAS of $\beta$-amyloid load (Amyloid) using individual-level ROS/MAP data and imputation models by **DPR (A)** and **PrediXcan (B)**.

**Figure S** 5

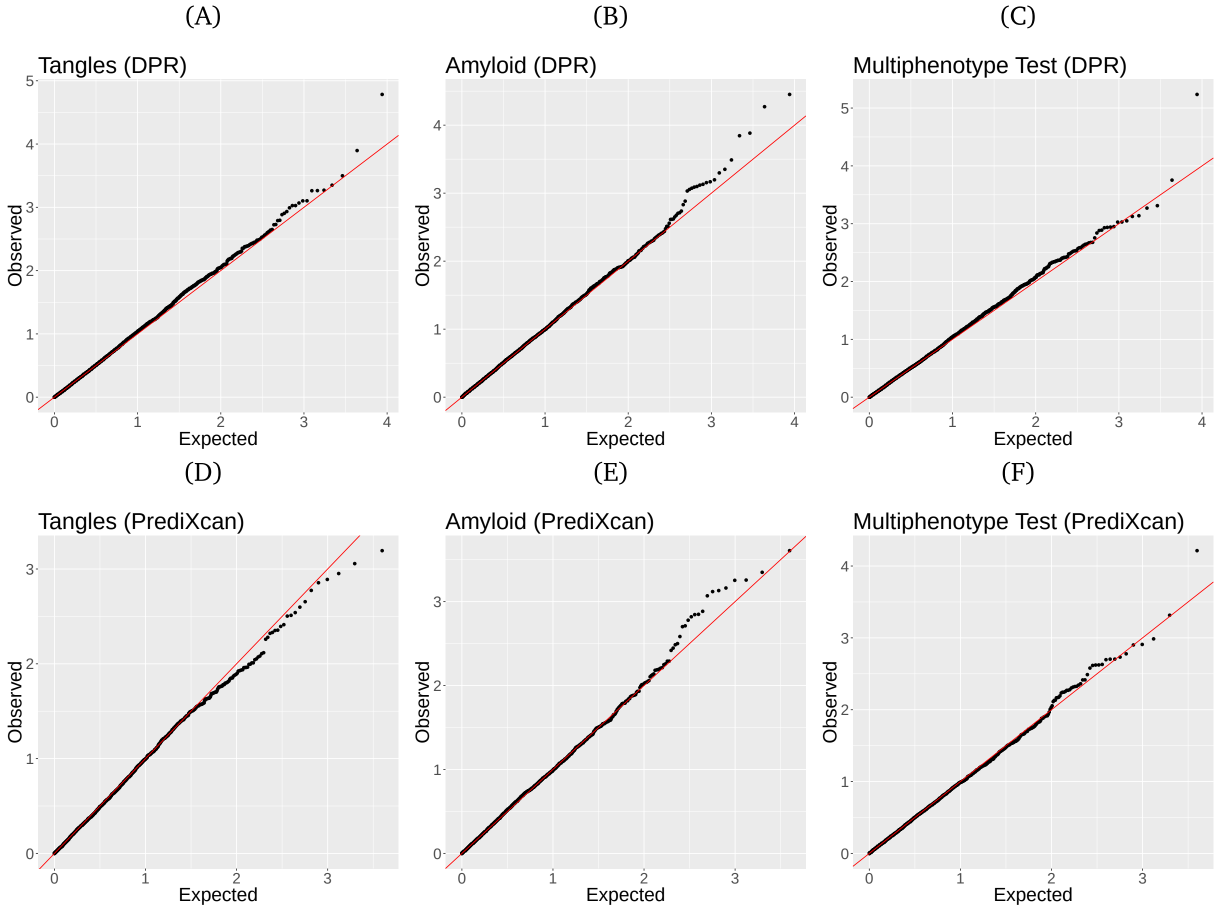


Quantile-Quantile (QQ) plots for univariate and multivariate TWAS of Tangles and Amyloid using individual-level ROS/MAP data and imputation models by **DPR (A, B, C)** and **PrediXcan (D, E, F)**.

**Figure S** 6

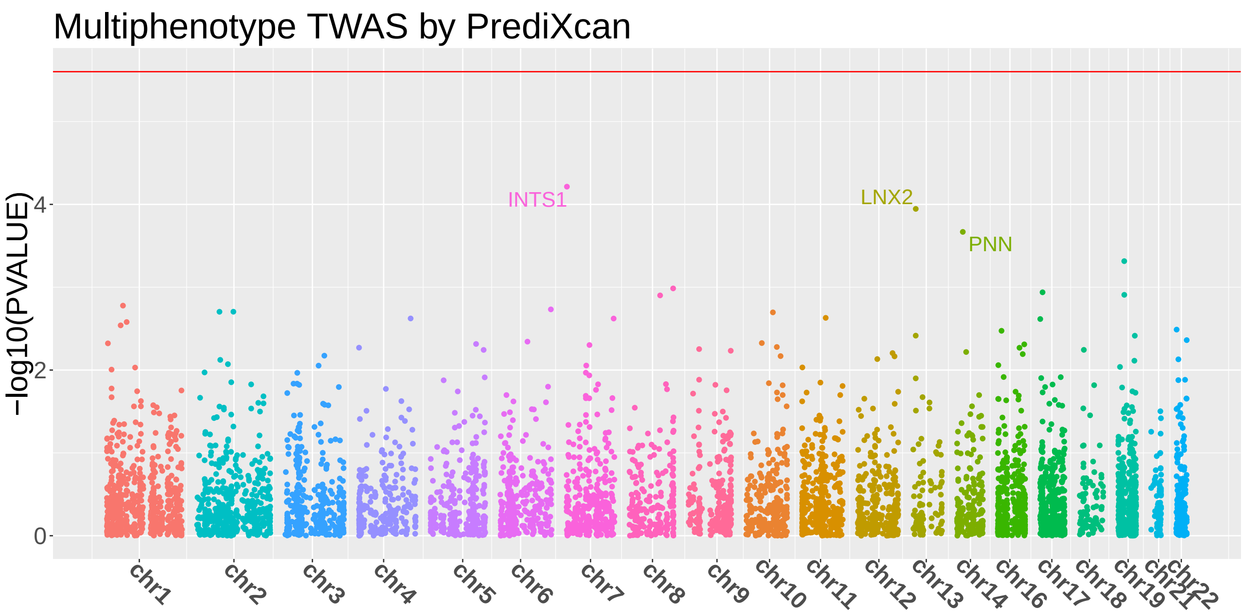


Manhattan plot for multiphenotype TWAS (with both Tangles and Amyloid) using individual-level ROS/MAP data by **PrediXcan**.
